# Structural Basis for Antiarrhythmic Drug Interactions with the Human Cardiac Sodium Channel

**DOI:** 10.1101/430934

**Authors:** Phuong T. Nguyen, Kevin R. DeMarco, Igor Vorobyov, Colleen E. Clancy, Vladimir Yarov-Yarovoy

## Abstract

The human voltage-gated sodium channel, hNa_v_1.5, is responsible for the rapid upstroke of the cardiac action potential and is target for antiarrhythmic therapy. Despite the clinical relevance of hNa_v_1.5 targeting drugs, structure-based molecular mechanisms of promising or problematic drugs have not been investigated at atomic scale to inform drug design. Here, we used Rosetta structural modeling and docking as well as molecular dynamics simulations to study the interactions of antiarrhythmic and local anesthetic drugs with hNav1.5. These calculations revealed several key drug binding sites formed within the pore lumen that can simultaneously accommodate up to two drug molecules. Molecular dynamics simulations identified a hydrophilic access pathway through the intracellular gate and a hydrophobic access pathway through a fenestration between domains III and IV. Our results advance the understanding of molecular mechanisms of antiarrhythmic and local anesthetic drug interactions with hNa_v_1.5 and will be useful for rational design of novel therapeutics.

## Introduction

Voltage-gated sodium channels (Na_V_) are transmembrane proteins that give rise to action potential generation and propagation in excitable cells. There are nine human Na_V_ (hNa_V_) channel subtypes expressed in neuronal, cardiac, and muscle cells (Catterall et al., 2005). The cardiac Na_V_ channel (Na_V_1.5) plays a central role in congenital and acquired cardiac arrhythmias and has been an important target for antiarrhythmic drug development (Chandra et al., 1999; Chen-Izu et al., 2015; DeMarco & Clancy, 2016; Dumaine & Kirsch, 1998; Fredj, Lindegger, et al., 2006; Moreno et al., 2011). Nevertheless, longstanding failures in drug treatment of heart rhythm disturbances and many other syndromes, which stem from a persistent failure to predict the effective or harmful action of drugs. For example, the CAST (“Preliminary report: effect of encainide and flecainide on mortality in a randomized trial of arrhythmia suppression after myocardial infarction. The Cardiac Arrhythmia Suppression Trial (CAST) Investigators,” 1989) and SWORD (Waldo et al., 1996) clinical trials showed that common antiarrhythmic drugs, such as encainide and flecainide, increased mortality and risk of sudden cardiac death in patients. Thirty years later, there is still no effective preclinical methodology to differentiate useful or potentially harmful drugs at the molecular level. In order to begin to develop and screen novel drugs to reveal the mechanisms of drug failure or efficacy for treatment of cardiovascular and other disorders (and to minimize side effects), a mechanistic understanding of drug interactions with Na_V_ channels at the atomic scale is needed.

Na_V_ channels respond dynamically to changes in cell membrane voltage and adopt distinct conformational states: open (conducting), closed (non-conducting) and inactivated (non-conducting). Na_V_ channels contain four homologous domains (DI-DIV), with each domain consisting of a voltage-sensing domain (VSD) containing transmembrane segments S1-S4 and a pore domain (PD) containing transmembrane segments S5 and S6 connected by a loop region with P1 and P2 helices forming selectivity filter (SF). Each VSD senses changes in membrane potential that leads to movement of its S4 segment which can, in turn, trigger channel activation (pore opening) or channel deactivation (pore closing) at the intracellular gate. The intracellular linker between domains DIII and DIV contains a hydrophobic isoleucine-phenylalanine-methionine (IFM) motif, which contributes to fast inactivation gating mechanism, resulting in rapid termination of Na^+^ conduction subsequent to the channel opening (Pan et al., 2018; Rohl et al., 1999; Shen et al., 2017; Vassilev et al., 1988; West et al., 1992; Yan et al., 2017). This inactivation process plays critical roles in Na_V_ channel function and drug binding (Catterall, 2014; Hille, 2001).

Gating and conduction in Na_V_ channels can also be modulated by drugs in a state-dependent manner (Hille, 1977; Hondeghem & Katzung, 1977). Inhibition of *I*_Na_ in a closed state is representative of a low affinity tonic block by neutral drugs accessing the Na_V_ receptor site through a hydrophobic pathway through the cell membrane (Buyan et al., 2018; Hille, 1977). However, many drugs that block *I*_Na_ access the Na_V_ receptor site through the intracellular hydrophilic pathway (Hille, 1977), and have a greater propensity for binding to the channel in open and inactivated states. In cardiac cells, drugs that exhibit slow unbinding kinetics during increased cell pacing can lead to use-dependent block (UDB), which has been shown to be potentially proarrhythmic (Moreno et al., 2011; Starmer et al., 1984). For this reason, investigations into the molecular determinants of the state dependence of drug binding to the open and inactivated states of Na_V_ channels is important for understanding what makes a certain class of drugs that target Nav channels safe, and others potentially proarrhythmic. Some of these drugs are commonly used as local anesthetics due to their action on neuronal Na_V_ channels, and thus their cardiac safety is of paramount importance (Reiz & Nath, 1986).

Forty years ago, Hille proposed two distinct access pathways for local anesthetics to the central binding site; the hydrophobic pathway through the membrane, and the hydrophilic pathway through the intracellular gate (Hille, 1977). Many antiarrhythmic and local anesthetic drugs are weak bases that exist in equilibrium between both neutral and charged forms at physiological pH. Neutral drugs may access the pore lumen binding site through both hydrophobic and hydrophilic pathways (Boiteux, Vorobyov, French, et al., 2014), but charged drugs are much more likely to access the pore binding site through the hydrophilic pathway, due to a large energetic penalty for traversing a lipid membrane (DeMarco et al., 2018). Extensive electrophysiological and site-directed mutagenesis experiments have identified a key receptor site for antiarrhythmic and local anesthetic drugs within the eukaryotic Na_V_ channel pore lumen (Ragsdale et al., 1994, 1996; Yarov-Yarovoy et al., 2001; Yarov-Yarovoy et al., 2002). Mutations of two conserved aromatic residues in the domain IV S6 (DIVS6) segment of Na_V_ channels, F1760 and Y1767 (hNa_v_1.5 numbering) significantly reduce antiarrhythmic and local anesthetic drug binding (Ragsdale et al., 1994, 1996). Other key residues for drug binding within the pore lumen have been identified in DIS6 and DIIIS6 segments (Yarov-Yarovoy et al., 2001; Yarov-Yarovoy et al., 2002). In addition, mutations within the Na_V_ channel selectivity filter region can affect drug binding, either through enhancement of slow inactivation or formation of alternative access pathway (P. J. Lee et al., 2001; Sunami et al., 1997; Tsang et al., 2005).

Structural studies have advanced our structural understanding of Na_V_ channel - drug interaction mechanisms. The first crystal structure of the bacterial Na_V_ channel Na_V_Ab revealed open fenestrations within the pore-forming domain (Payandeh et al., 2012; Payandeh et al., 2011), which supported the hypothesis that drugs can access the binding site within the pore lumen through the hydrophobic pathway. Crystal structures of Na_V_Ms and Ca_V_Ab channels have been determined with drugs bound near the fenestration regions or in the pore lumen, suggesting the possibility of similar drug binding receptor sites in eukaryotic Na_V_ channels (Bagneris et al., 2014; L. Tang et al., 2016). The first high-resolution structures of eukaryotic Na_V_ channels have recently been resolved using cryo-electron microscopy (cryoEM). The *American cockroach* Na_V_PaS channel structures have been solved in a closed state (Shen et al., 2018; Shen et al., 2017) and *electric eel* Na_V_1.4 channel structure has been solved in a partially open and presumably inactivated state (Yan et al., 2017). These structures have unlocked new opportunities to study drug interactions with eukaryotic Na_V_ channels at the atomic scale.

The Rosetta computational modeling software (Alford et al., 2017; Bender et al., 2016; Rohl et al., 2004; Simons et al., 1999) has been used to study conformational changes in Na_V_, voltage-gated potassium (K_v_), voltage-gated calcium (Ca_V_), and TRPV1 channels (Decaen et al., 2011; DeCaen et al., 2009; DeCaen et al., 2008; P. T. Nguyen et al., 2017; Pathak et al., 2007; Tuluc et al., 2016; Vargas et al., 2012; Yang et al., 2018; Yarov-Yarovoy et al., 2006; Yarov-Yarovoy et al., 2012) and peptide toxin interactions with Na_V_, K_v_, and TRPV1 channels (Catterall et al., 2007; Cestele et al., 2006; Gupta et al., 2015; Kimball et al., 2016, 2018; Kimball et al., 2017; P. T. Nguyen et al., 2015; P.T. Nguyen et al., 2014; C. Tang et al., 2017; Tilley et al., 2014; J. Wang et al., 2011; S. Yang et al., 2015; Zhang et al., 2011, 2012). RosettaLigand flexible docking (DeLuca et al., 2015) has been used to study small molecule interactions with Nav, TRPV1 and calcium-activated K^+^ channels (H. M. Nguyen et al., 2017; P. T. Nguyen et al., 2018; Yang et al., 2016; F. Yang et al., 2015). Molecular dynamics (MD) simulations have previously revealed drug binding and access to bacterial Na_V_ channels (Barber et al., 2014; Boiteux, Vorobyov, French, et al., 2014; Corry et al., 2014; Martin & Corry, 2014). Molecular docking of antiarrhythmic, local anesthetic, and anticonvulsant drugs with homology models of a eukaryotic Na_V_1.4 channel based on bacterial Na_V_Ms channel in an open state, has recently revealed electroneutral and cationic drug interactions with the phenylalanine in the DIVS6 segment (F1760 in human Na_V_1.5) and selectivity filter region (Tikhonov & Zhorov, 2017). Differences in binding of neutral and charged local anesthetics have been recently studied using the bacterial Na_V_Ms channel in an open state and eukaryotic Na_V_PaS channel in a closed state (Buyan et al., 2018). Structural, experimental, and modeling studies have all provided a better understanding of drug interactions with bacterial Na_V_ channels and models of eukaryotic Na_V_ channels in open or closed states. However, atomistic details remain elusive for antiarrhythmic and local anesthetic drug access pathways, specific binding sites, and stoichiometry of binding to eukaryotic Na_V_ channels in an inactivated state, which forms high affinity drug binding site (Carnevale, 2018).

In this study, we used Rosetta to build a model of the human Na_V_1.5 (hNa_v_1.5) channel in a partially open and presumably inactivated state based on the cryo-EM structure of the electric eel Na_V_1.4 channel and conducted a docking study to investigate the interactions of antiarrhythmic and local anesthetic drugs - lidocaine, QX-314, etidocaine, flecainide, and ranolazine - with hNav1.5. The results revealed that both antiarrhythmic and local anesthetic drugs share a receptor site formed by the S6 segments from domains III and IV. Multi-microsecond unbiased MD simulations of neutral lidocaine interacting with hNa_v_1.5 using the Anton 2 supercomputer revealed a hydrophilic access pathway through the intracellular gate, and a novel hydrophobic access pathway through a fenestration between domains III and IV. Distinct binding sites were identified in the pore region for both neutral and charged lidocaine. And we observed that the channel can accommodate up to two lidocaine molecules binding at the same time. Our results reveal the high-resolution structural determinants of drug block of hNa_v_1.5 in an inactivated state. They also serve as initial steps toward linking of structural determinants of channel - drug interactions to the modification of hNa_v_1.5 function.

## Results and discussion

### A structural model of the human Na_V_1.5 channel based on electric eel Nav1.4 channel structure

To study the state-dependent molecular mechanisms of high affinity binding of antiarrhythmic and local anesthetic drugs to human Na_V_ channels at the atomic scale, high-resolution structures of eukaryotic Na_V_ channels in open and inactivated states are needed. The cryoEM structure of the electric eel Na_V_1.4 (eeNa_V_1.4) channel in a partially open and presumably inactivated state (PDB ID: 5XSY) (Yan et al., 2017) provides atomic accuracy structural template for modeling of human Na_V_ channels. The sequence identity between hNa_v_1.5 and eeNa_V_1.4 is ~84% in the pore-forming transmembrane region (Figure 1 - figure supplement 1), which is within an atomic level accuracy homology modeling range (Koehl & Levitt, 1999; Marti-Renom et al., 2000), allowing us to generate accurate model of hNa_v_1.5 in a partially open and presumably inactivated state.

The human Nav1.4 (hNav1.4) structure was just published in September of 2018 (Pan et al., 2018), when this study was already completed. The sequence identity in the between hNa_v_1.5 and hNa_V_1.4 is only slightly higher (~87%) than sequence identity between hNa_v_1.5 and eeNa_V_1.4 (~84%) over the pore-forming transmembrane region, which suggests that eeNav1.4 and hNav1.4 structures are within the same range of accuracy for modeling of human Nav channels. The overall root mean square deviation (RMSD) between hNav1.4 and eeNav1.4 structures is less than 1 Å (Pan et al., 2018) and RMSD over the pore-forming domain segments S5 and S6 and P1- and P2-helices is less than 0.7 Å, which suggests very similar conformations of the pore-forming domain structure - the main focus of this study.

The eeNa_V_1.4 structure has the following distinct structural features: 1) a partially open intracellular gate in the PD; 2) an activated state of domain III and IV VSDs; 3) an inactivation gate (“IFM” motif in domain III-IV linker) bound between S4-S5 linkers in domains III and IV and DIVS6 segment (Yan et al., 2017). Based on these observations, the eeNa_V_1.4 structure potentially represents a partially open and presumably inactivated state, which has high affinity for antiarrhythmic and local anesthetic drugs (Ragsdale et al., 1994, 1996). We used the Rosetta structural modeling software (Alford et al., 2017; Bender et al., 2016; Rohl et al., 2004) with the eeNa_V_1.4 channel structure as a template to build a homology model of hNa_v_1.5 channel in a partially-open-inactivated state as described in Materials and Methods (Figure 1).

**Figure 1.**
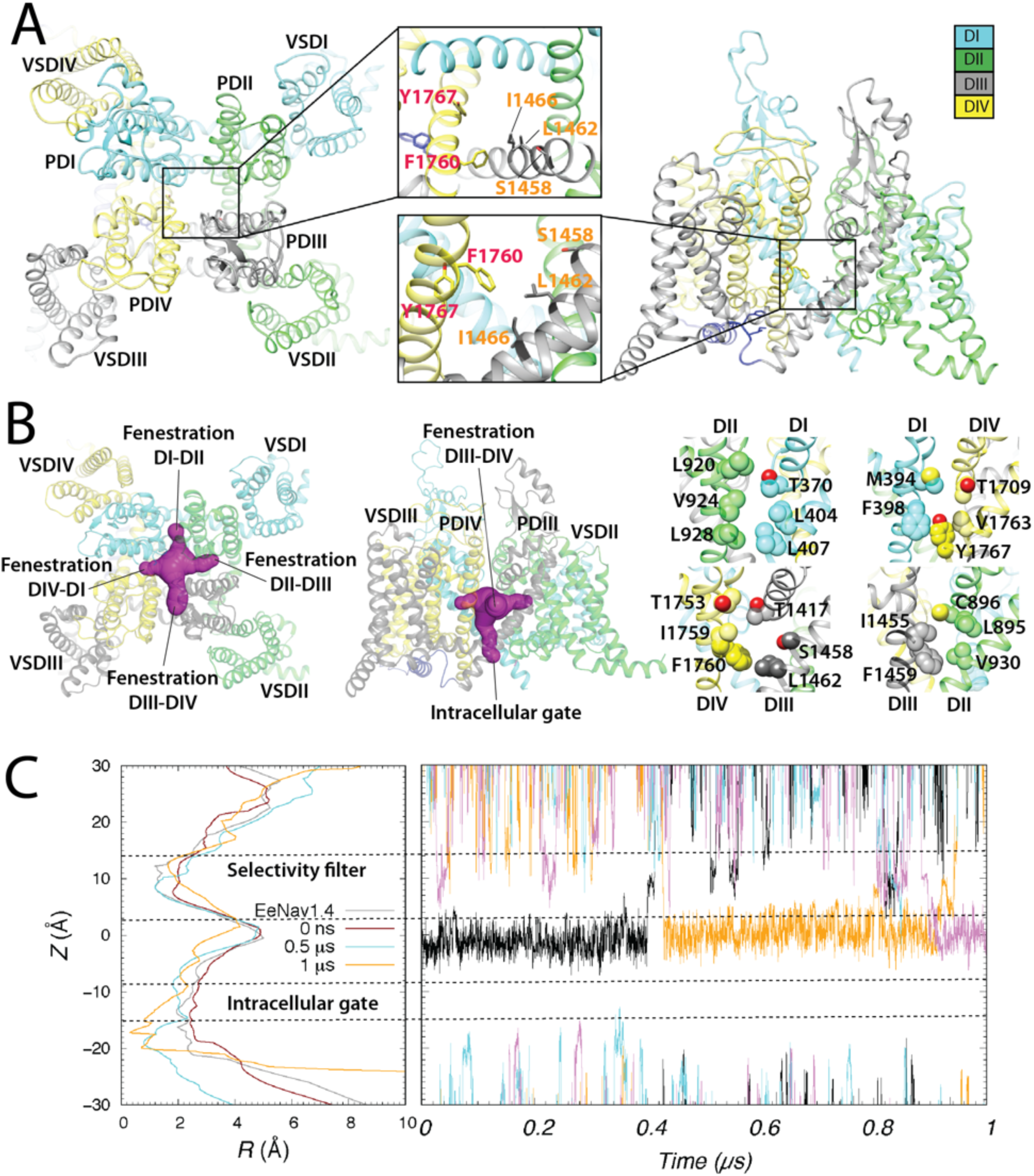
Rosetta model of the hNa_V_1.5 channel. (A) Extracellular (left panel) and transmembrane (right panel) views of the hNa_v_1.5 model shown in ribbon representation. Insets - zoom-in views of putative drug binding residues within hNav1.5 pore lumen. Each domain is colored inDIVidually and labeled. In the insets, DIII residues are labeled orange, whereas DIV residues are labelled red. (B) Extracellular (left) and transmembrane (center and right) views of all four hNav1.5 fenestrations using molecular surface representation (shown in purple in the left and center panels). In the right panels, fenestration-facing residue side chains are labelled and shown in space-filling representations using corresponding domain colors, with O atoms shown in red. (C) *Left panel*, hNav1.5 pore lumen radius (*R*) profile changes during molecular dynamic simulation at time zero (colored red), at 0.5 μs (colored cyan), and at 1 μs (colored orange). A pore lumen *R* profile for a cryoEM eeNa_V_1.4 structure is also shown in gray for comparison. *Right panel*, Sodium ion trajectories within the pore-forming domain during a 1 μs molecular dynamic simulation of hNav1.5.

Key amino acid residues forming the putative antiarrhythmic and local anesthetic drug binding site in DIIIS6 and DIVS6 segments (Ragsdale et al., 1994, 1996; Yarov-Yarovoy et al., 2001; Yarov-Yarovoy et al., 2002) are identical between hNa_v_1.5 and eeNa_V_1.4 (Figure 1 - figure supplement 1). For example, F1760 and Y1767 in the DIVS6 segment in hNa_v_1.5 (Figure 1A) are F1555 and Y1562 in eeNa_V_1.4, respectively. Moreover, L1462 and I1466 in the DIIIS6 segment in hNa_v_1.5 (Figure 1A) are L1256 and I1260 in eeNa_V_1.4, respectively. I1756 in the DIVS6 segment in hNa_v_1.5 is also identical in eeNa_V_1.4 (I1551) and forms part of the drug access pathway at the fenestration between the DIIIS6 and DIVS6 segments (see Figure 1 - figure supplement 1) (Ragsdale et al., 1994). However, another key amino acid residue in the drug access pathway at the fenestration between DIIIS6 and DIVS6 segments (Qu et al., 1995) is different between hNa_v_1.5 and eeNay1.4: T1753 in the DIVS6 segment of hNay1.5 is C1548 in eeNay1.4 (see Figure 1B, Figure 1 - figure supplement 1, and Figure 1 - figure supplement 1). Notably, T1753 is facing L1413 in the P1-helix of DIII, which is a unique residue in the fenestration between the DIIIS6 and DIVS6 segments because all other Na_V_ channel domains have a Phenylalanine at the corresponding position (see Figure 1 - figure supplement 1 and Figure 1 - figure supplement 1). These unique structural features of the fenestration between the DIIIS6 and DIVS6 segments will be relevant for the MD simulations of the lidocaine access pathway discussed below.

To determine whether the hNa_v_1.5 channel model represents a conductive or non-conductive open state, we performed molecular dynamics (MD) simulations of the hNa_v_1.5 model as described in Materials and Methods. The Rosetta hNav1.5 model and the eeNa_V_1.4 structure both have a ~2.5 Å pore radius within the intracellular gate region (Figure 1C, left panel) (Yan et al., 2017). During the MD simulation of the hNav1.5 model, the intracellular gate radius decreased from ~2.5 Å at the start of the simulation to ~2.0 Å after 0.5 μs and then to ~1.0 - 2.0 Å after 1 μs (Figure 1C). While we observed several Na^+^ ions passing up and down between the selectivity filter region and the pore lumen, we did not detect any Na^+^ ions passing through the intracellular gate of the pore during the 1 μs simulation (Figure 1C, right panel). Based on these results, we assume our hNa_v_1.5 model to be in a non-conductive inactivated state.

### Modeling of antiarrhythmic and local anesthetic drugs interaction with human Na_V_1.5 channel using RosettaLigand

To study high affinity binding of antiarrhythmic and local anesthetic drugs to the hNa_v_1.5 pore in the non-conductive inactivated state at atomic scale, we used RosettaLigand (Bender et al., 2016; Davis & Baker, 2009; DeLuca et al., 2015; Lemmon & Meiler, 2012; Meiler & Baker, 2006) as described in Materials and Methods.

*Lidocaine* is a local anesthetic and class Ib antiarrhythmic drug used for the treatment of ventricular arrhythmias (Singh, 1997). Experimental data suggest that phenylalanine and tyrosine residues in the DIVS6 segment of mammalian Na_V_ channels (F1760 and Y1767 in hNa_v_1.5) play a key role in antiarrhythmic and local anesthetic drug binding (Ragsdale et al., 1996). The most frequently sampled lowest binding energy RosettaLigand models of neutral or charged lidocaine interacting with hNav1.5 indicate that the region above F1760 in the DIVS6 segment forms a “hot spot” for lidocaine binding (Figure 2A and B and Figure 2 - figure supplement 2 and 2). This “hot spot” extends from the fenestration between the DIIIS6 and DIVS6 segments into the pockets under the selectivity filter region in DIII and DIV. The tertiary amine group of neutral and charged lidocaine is positioned above F1760 (Figure 2A and B). The phenyl ring of neutral and charged lidocaine is observed in multiple different orientations near F1760 (Figure 2A and B and Figure 2 - figure supplement 2 and 2). We observed only one neutral and one charged lidocaine pose among the lowest energy models near Y1767 (Figure 2 - figure supplement 2 and 2), potentially reflecting a lower affinity binding site near this residue and in agreement with a weaker impact of Y1767 mutations on drug binding compared to F1760 mutations (Ragsdale et al., 1994, 1996). Experimental data suggest that leucine and isoleucine residues in the DIIIS6 segment of mammalian Na_V_ channels (L1462 and I1466 in hNa_v_1.5) also form receptor site for antiarrhythmic and local anesthetic drug binding (Nau et al., 2003; Yarov-Yarovoy et al., 2001). The L1462 residue is positioned near F1760 in our model (Figure 2A and B). However, I1466 is not in direct contact with lidocaine in any of top neutral and charged lidocaine models, suggesting an allosteric effect of mutations at this position on drug binding.

**Figure 2.**
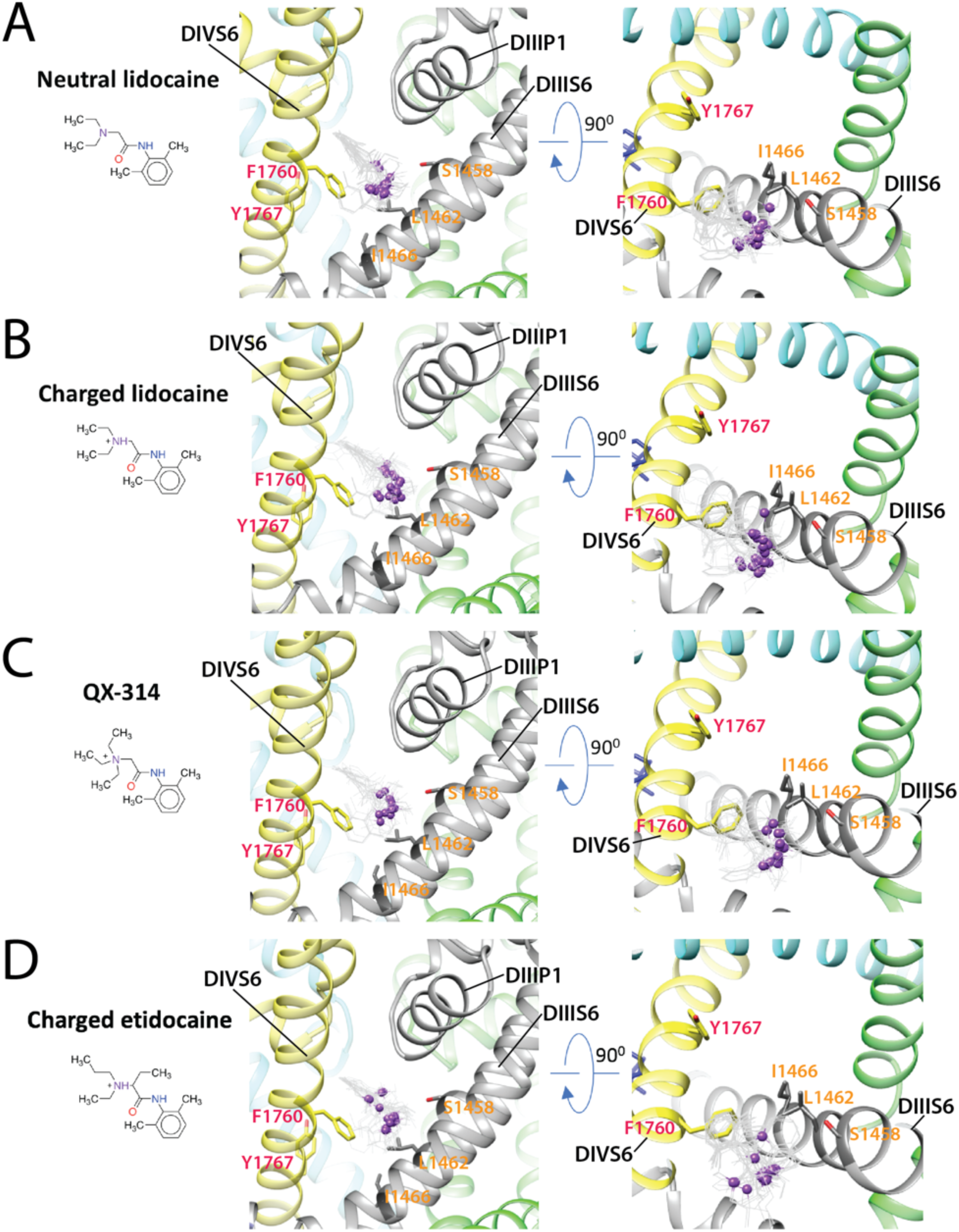
Rosetta models of hNa_v_1.5 channel interaction with antiarrhythmic and local anesthetic drugs. Close up transmembrane (left panels) and extracellular (right panels) views of hNa_v_1.5 interactions with: (A) neutral lidocaine; (B) charged lidocaine; (C) QX-314; (D) charged etidocaine. Drug molecules are shown in the wireframe representations with basic N atoms depicted as purple balls. hNa_v_1.5 domain I is colored in blue, domain II is colored in green, domain III is colored in gray, domain IV is colored in yellow. Side chains of key residues forming the receptor site in DIIIS6 and DIVS6 segments are shown in stick representation and labeled in orange and red, respectively.

To validate the robustness of the RosettaLigand prediction of the “hot spot” for lidocaine binding, we explored modeling of two well-studied lidocaine variants - QX-314 and etidocaine.

*QX-314* is a permanently charged derivative of lidocaine with a quaternary ammonium group. The most frequently sampled lowest binding energy RosettaLigand models of QX-314 interacting with hNav1.5 indicate that the region above F1760 in the DIVS6 segment forms a “hot spot” for QX-314 binding (Figure 2C and Figure 2 - figure supplement 2), which is similar to the “hot spot” observed in our lidocaine - hNav1.5 models. The ammonium group of QX-314 is positioned above F1760 (Figure 2C). The phenyl ring of QX-314 is observed in multiple different orientations near F1760 (Figure 2C and Figure 2 - figure supplement 2).

*Etidocaine* is a local anesthetic drug that was used in the first experimental study by the Catterall group that identified key residues of the receptor site for state-dependent block in both the DIVS6 segment (F1760 and Y1767 in hNa_v_1.5) (Ragsdale et al., 1994) and the DIIIS6 segment (L1462 and I1466 in hNa_v_1.5) (Yarov-Yarovoy et al., 2001). The most frequently sampled lowest binding energy RosettaLigand models of charged etidocaine show the molecule binding above F1760 in the DIVS6 segment (Figure 2D and Figure 2 - figure supplement 2), which is similar to the “hot spot” observed in our lidocaine and QX-314 - hNav1.5 models. The ammonium group of etidocaine is positioned above and near F1760 (Figure 2D). The phenyl ring of etidocaine is observed in multiple different orientations near F1760 (Figure 2D and Figure 2 - figure supplement 2).

*Flecainide* is a class 1c antiarrhythmic drug used to prevent and treat tachyarrhythmias, which also may have unpredictable proarrhythmic effects (Anderson et al., 1984; Benhorin et al., 2000; Holmes & Heel, 1985; Liu et al., 2003; Liu et al., 2002). Experimental data suggest that flecainide preferentially binds to Na_V_ channels in an open state and that phenylalanine and tyrosine residues in the DIVS6 segment (F1760 and Y1767 in hNa_v_1.5) play an important role in its binding (Liu et al., 2003; Liu et al., 2002; Ragsdale et al., 1996; G. K. Wang et al., 2003). The most frequently sampled lowest binding energy RosettaLigand models of flecainide in hNav1.5 are consistent with the other drugs in that the region above F1760 in the DIVS6 segment also forms a “hot spot” for flecainide binding (Figure 3A and Figure 3 - figure supplement 1). However, the larger and branched structure of flecainide compared to lidocaine, etidocaine, and QX-314 results in a greater surface area of interaction that spans from the fenestration region between the DIII and DIV to the ion conduction pathway under the selectivity filter region (Figure 3A).

**Figure 3.**
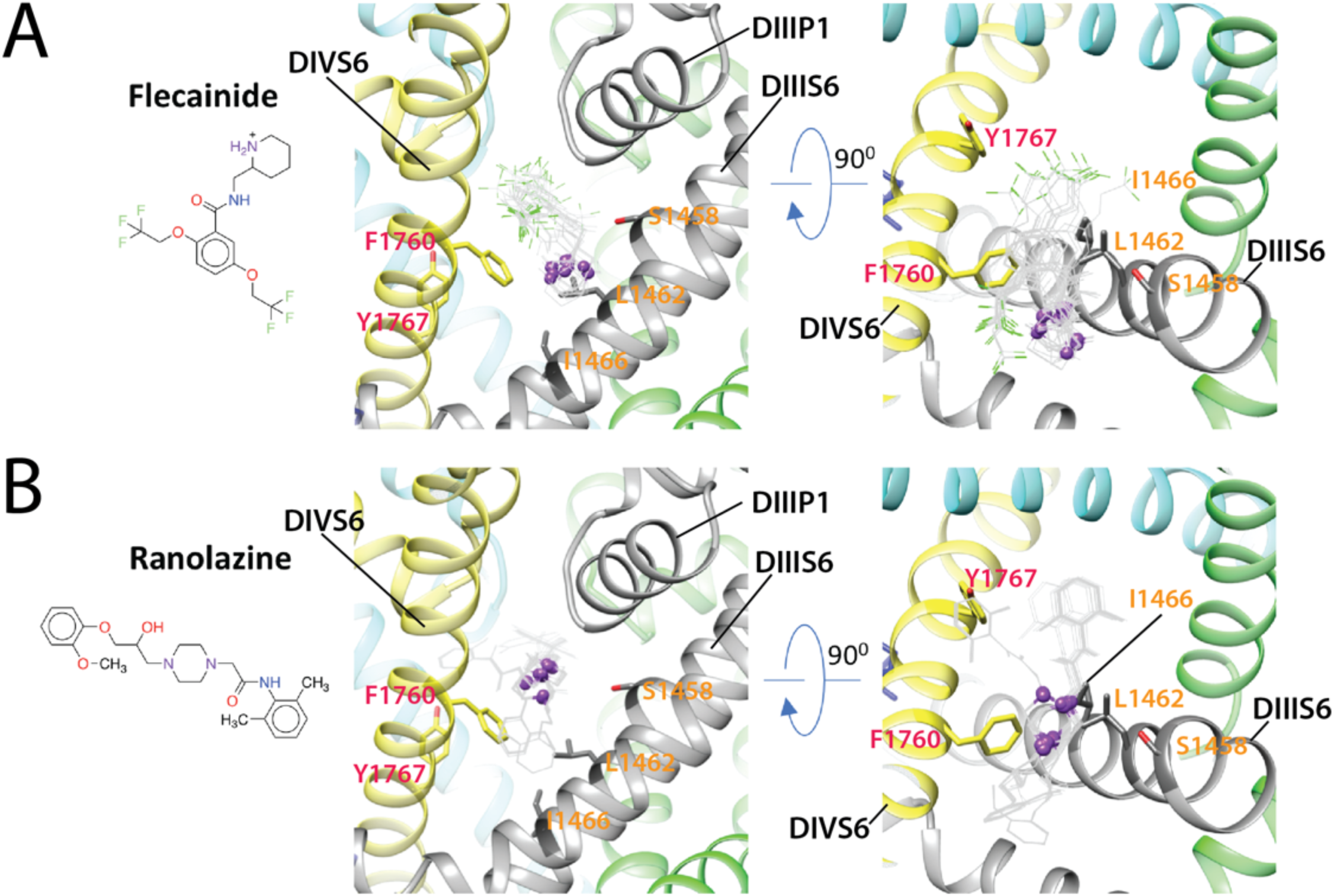
Rosetta models of hNa_v_1.5 channel interaction with antiarrhythmic and local anesthetic drugs. Close up transmembrane (left panel) and extracellular (right panel) view of hNa_v_1.5 interactions with (A) flecainide; (B) ranolazine. Drug molecules are shown in the wireframe representations with flecainide F atoms colored in green and basic N atoms of both drugs depicted as purple balls. hNa_v_1.5 domain I is colored in blue, domain II is colored in green, domain III is colored in gray, domain IV is colored in yellow. Side chains of key residues forming the receptor site in DIIIS6 and DIVS6 are shown in stick representation and labeled in orange and red, respectively.

*Ranolazine* is an anti-anginal drug that inhibits late Na_V_ current. Experimental data suggest that ranolazine binds to Na_V_ channels in an open state and that phenylalanine in the DIVS6 segment (F1760 in hNa_v_1.5) plays key role in its binding (Fredj, Sampson, et al., 2006; G. K. Wang et al., 2008). The most frequently sampled lowest binding energy RosettaLigand models of ranolazine show that the same region above F1760 in the DIVS6 segment forms the “hot spot” for ranolazine binding (Figure 3B and Figure 3 - figure supplement 2). Ranolazine has a flexible linear rather than branched structure and interacts via the same modality as flecainide over a larger surface area that spans from the fenestration region between the DIII and DIV to the ion conduction pathway under the selectivity filter region (Figure 3B).

Overall, the RosettaLigand docking results suggest that the region above F1760 in DIVS6 forms a “hot spot” for binding of antiarrhythmic and local anesthetic drugs and includes the interface between the DIIIS6 and DIVS6 segments and the pocket under the selectivity filter region in DIII and DIV. The key role of F1760 in hNa_v_1.5 and the equivalent phenylalanine residue in other Na_V_ channels agrees with experimental data for multiple antiarrhythmic and local anesthetic drugs (Fredj, Sampson, et al., 2006; Liu et al., 2003; Liu et al., 2002; Ragsdale et al., 1994, 1996; G. K. Wang et al., 2008; G. K. Wang et al., 2003). Positioning of the drugs between the DIIIS6 and DIVS6 segments in our models is in agreement with the Chanda Lab structural hypothesis that local anesthetics may act as a “wedge” to stabilize primarily VSDIII and partially VSDIV in activated states (Muroi & Chanda, 2009). The position of the drugs under the selectivity filter region in DIII and DIV is notable with respect to several mutations in this region that have been shown to significantly affect the slow inactivation of Na_V_ channels (Balser et al., 1996; Kambouris et al., 1998; Ong et al., 2000; Todt et al., 1999). We hypothesize that upon binding above F1760 in DIVS6 and under the selectivity filter region in DIII and DIV the antiarrhythmic and local anesthetic drugs may induce conformational changes that may enhance slow inactivation of Na_V_ channels in agreement with experimental data (Chen et al., 2000; Fukuda et al., 2005). We also propose that since the antiarrhythmic drugs ranolazine and flecainide have more extensive interactions with the channel compared to lidocaine and its derivatives in our models (see Figures 2 and 3), their effect on channel gating might be more prominent as well. In fact, our recent multi-scale kinetic modeling and experimental study examined lidocaine and flecainide interactions with Na_V_1.5 and their consequence on pro-arrhythmia proclivities (Moreno et al., 2011). We found, for example, that cardiac-safe lidocaine has faster channel unbinding kinetics, resulting in more facile recovery of channels from drug blockade, and lower incidence of reentrant arrhythmias at a cardiac tissue and a whole heart level compared to flecainide.

### Neutral and charged lidocaine partitioning into the membrane

The molecular docking calculations, described above, provided us with atomistic structural models of convergent binding poses of several anti-arrhythmic and local anesthetic drugs in the hNa_v_1.5 pore. However, static molecular models cannot tell us how a drug accesses the binding site and whether such drug - protein interactions are long-lived or transient. Such information can be provided by atomistic molecular dynamics (MD) simulations of a channel embedded in a hydrated lipid membrane with one or multiple drug molecules present. To perform such simulations, we need accurate atomic-resolution structural models, called empirical force fields, for all the system components. For this study, we used biomolecular and generalized all-atom CHARMM force fields, which were previously utilized by our and other groups to study bacterial Na_V_ channel conduction and drug binding (Boiteux, Vorobyov, & Allen, 2014; Boiteux, Vorobyov, French, et al., 2014; Chakrabarti et al., 2013; Corry & Thomas, 2012; Lenaeus et al., 2017; Martin et al., 2014).

We focused the MD simulations on hNav1.5 interactions with charged and neutral forms of lidocaine. This widely used antiarrhythmic and local anesthetic drug was chosen for our exploratory MD study because molecular docking calculations and previous experimental data indicate that it shares the same binding site as larger Na_V_1.5 blockers such as flecainide and ranolazine. Our previous MD simulation study of drug - bacterial Nav channel interactions suggested that we can more efficiently predict entry and egress pathways for a smaller drug, like the local anesthetic benzocaine, compared to the larger anti-epileptic drug phenytoin (Boiteux, vorobyov, French, et al., 2014). Indeed, experimental data indicate that lidocaine has faster Na_V_1.5 association and dissociation kinetics than the larger flecainide (Moreno et al., 2011). Moreover, in aqueous solution lidocaine exists as a mixture with a substantial fractions of both charged (~78% at pH=7.4) and neutral form (~22% at pH=7.4) which have different membrane permeabilities and can interact with the ion channels via distinct pathways, as was discussed above. Previous experimental and simulation studies suggested that charged and neutral forms of lidocaine differently affect Na_V_ channel function (Buyan et al., 2018; Moreno et al., 2011; O’Leary & Chahine, 2018; Tikhonov & Zhorov, 2017). Therefore, in this study we have explored charged and neutral lidocaine - lipid membrane and Na_V_1.5 interactions via all-atom MD simulations. We developed force field parameters for charged and neutral lidocaine, because they are not available in the standard biomolecular (Huang & MacKerell, 2013; Klauda et al., 2010) or generalized CHARMM force field (CGENFF) (Vanommeslaeghe et al., 2010b). We used gas-phase quantum mechanical (QM) drug geometries, vibrational frequencies, dihedral angle profiles, dipole magnitude and direction as well as interactions with water in different orientations as reference values for the parameter development, as described in Appendix and illustrated in Figure 4 - figure supplement 1 and 2 and Tables S1-S3.

The derived parameters were validated by performing MD simulations of charged and neutral lidocaine partitioning across a 1-palmitoyl-2-oleoyl-phosphatidylcholine (POPC) lipid membrane and computing the water-membrane distribution coefficient log *D* = 1.25, which agrees favorably with the experimental value of 1.76 (Avdeef et al., 1998). Lidocaine free energy profiles, used to obtain our logD estimate using Eq. 2 below are shown in Figure 4 - figure supplement 3 and demonstrate that there is a higher barrier for charged vs. neutral lidocaine translocation across a lipid membrane in agreement with a previous study using different drug models (Buyan et al., 2018). However, contrary to ~5 kcal/mol free energy well at the membrane center for neutral lidocaine in that study (Buyan et al., 2018), our simulations predict an interfacial minimum of - 1.09 kcal/mol at |*z*| = 13 Å and a ~4.64 kcal/mol peak at the membrane center (Figure 4 - figure supplement 3). We also obtained even more favorable interfacial binding of -3.07 kcal/mol at |*z*| = 15 Å for charged lidocaine, which despite a larger peak of 6.58 kcal/mol at the membrane center leads to a more favorable membrane partitioning of this form (cf. partitioning coefficients for neutral and charged lidocaine forms, log*K_0_* = 0.12 and log*K_1_* = 1.35 respectively). We also used an approximation of Kramer’s transition rate theory to estimate the transition rates (Allen et al., 2003; Crouzy et al., 1994) of charged and neutral forms of lidocaine through a simulated POPC bilayer. We used the same approach as in our previous study (DeMarco et al., 2018) and for charged and neutral lidocaine computed their diffusion coefficients (Hummer, 2005) close to the membrane center using Hummer’s method, as well as the curvatures around the binding wells and peaks (i.e. free energy minima and maxima), estimated from second derivatives of second-order polynomial fits to the relevant portion of each respective free energy profile. Estimated transition rates through the membrane are 38.9s^−1^ for charged lidocaine and 21.1ms^−1^ for the neutral drug form, indicating three orders of magnitude faster crossing rate for the latter.

Since charged lidocaine is the dominant drug form at a physiological pH 7.4 (~78.4% based on its p*K*_a_ = 7.96) (Pless et al., 2011), we primarily expect the accumulation of charged drug at water-membrane interfaces, in agreement with recent solid NMR experiments (Weizenmann et al., 2012). However, deeper into the hydrophobic membrane core, neutral lidocaine is expected to be the more dominant form and should be able to translocate across a membrane more easily due to the substantially smaller barrier than its protonated counterpart (~6 kcal/mol vs. ~10 kcal/mol) (Figure 4 - figure supplement 3). This indicates that we need to study both charged and neutral lidocaine interactions with hNa_v_1.5 to assess hydrophobic (lipid-mediated access through channel fenestrations) and hydrophilic (water-mediated access through an intracellular gate) channel pore drug access pathways and understand molecular mechanisms of channel activity modulation.

### Molecular dynamics simulations reveal neutral lidocaine access pathways to the binding site via the intracellular gate and fenestration between domains III and IV

To explore the lidocaine access pathways to its binding site within the hNa_v_1.5 channels, we ran multi-microsecond MD simulations on the Anton 2 supercomputer (Shaw et al., 2014) with neutral or charged lidocaine, as described in Materials and Methods. The MD simulations of neutral lidocaine revealed that it can access its binding site within the Na_V_ channel pore lumen either through an opening formed by the intracellular gate (hydrophilic pathway) or through a path formed between the lipids, the P1-helix in DIII, the P2-helix in DIV, and the fenestration region between domains III and IV (hydrophobic pathway) (see Figure 4 and Supplemental Movies 1 and 2). The hydrophilic pathway is formed by the following residues at the intracellular gate (see sites I1 and I2 in Figure 4A and C): L404, I408, V412 (DIS6), L931, F934, L935, L938 (DIIS6), L1462, I1466, I1470 (DIIIS6), and V1764, Y1767, I1768, I1771 (DIVS6). Notably, all of the residues lining the intracellular gate in human Na_V_ channels are hydrophobic and highly conserved. The hydrophobic pathway between domains III and IV is formed by the following residues (see sites E1, E2, C1, and C2 in Figure 4A and 4C): L1338, L1342, W1345 (in DIIIS5), L1410, L1413, Q1414 (in P1-helix of DIII), L1462, F1465 (in DIIIS6), W1713, L1717, L1721 (in P2-helix of DIV), and I1749, T1753, I1756, I1757 (in DIVS6). Remarkably, lidocaine molecules that accessed the pore binding sites (C1, C2 sites) are not those partitioned from lipid membrane. Lidocaine accessed the fenestration between domains III and IV from the extracellular side by going through the cleft formed between P1-DIII and P2-DIV (E1, E2 sites). Furthermore, F1760 (in DIVS6) and L1462 (in DIIIS6) are the first residues that lidocaine encounters as it enters the pore lumen through the fenestration region - both of these residues are forming the “hot spot” for all the drugs simulated using RosettaLigand (see Figures 2 and 3). Moreover, neutral lidocaine was found to access the receptor site via the fenestration between domains III and IV, but not through the fenestrations between the other domains. We hypothesize that specific amino acid differences between the residues forming the fenestration between domains III and IV versus residues forming fenestrations between all other domains are preventing lidocaine from accessing the receptor site through the other fenestrations (Figure 1 - figure supplement 1 and Figure 1 - figure supplement 1).

**Figure 4.**
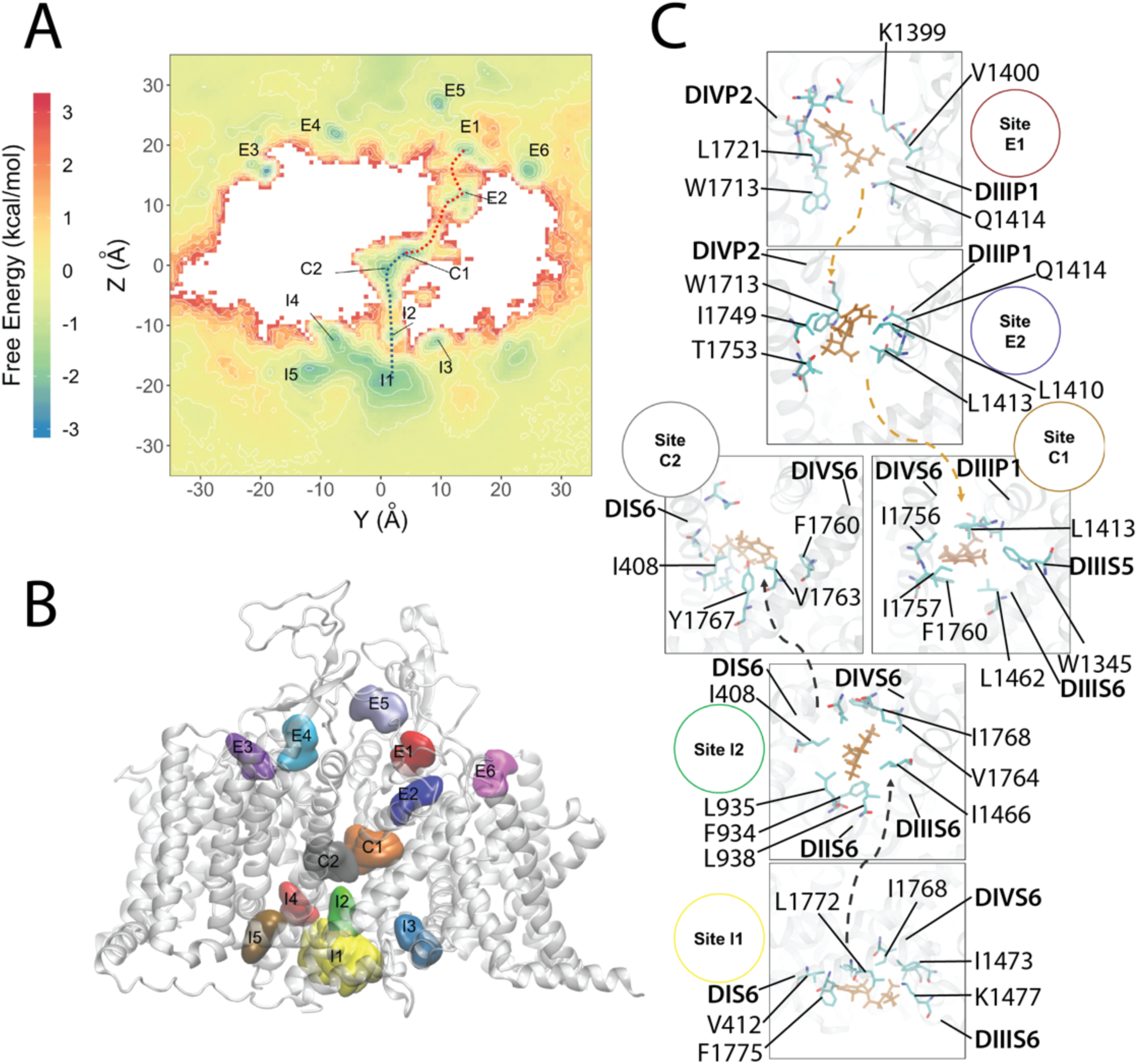
Molecular dynamics simulation of the hNa_v_1.5 channel interaction with neutral lidocaine reveals two drug access pathways. (A) Free energy surface of neutral lidocaine binding projected on the *Y-Z* plane (with Z corresponding to a transmembrane axis). Binding sites for neutral lidocaine, identified from free energy minima, are labeled as intracellular I1–5, channel pore C1–2, and extracellular E1–6. (B) Transmembrane view of the channel with neutral lidocaine binding sites represented as colored surfaces. Colors and sizes are for clarity, not actual binding properties. (C) Close-up view of binding sites forming the hydrophobic (orange arrows) and hydrophilic (gray arrows) binding pathways. Lidocaine molecules (orange) and interacting residues on the channel (cyan for C, red for O and blue for N) are shown using stick representation.

We found this observation of the hydrophobic pathway very intriguing. Although early work on local anesthetics and quaternary derivatives provided compelling evidence for a hydrophobic pathway as a result of drug partitioning into lipid membrane (Frazier et al., 1970; Hille, 1977; Narahashi et al., 1970; Strichartz, 1973), variants of different channel isoforms appeared to have a specific residue dependent external access pathway. Membrane-impermeant QX-314 was shown to block the cardiac isoform Na_V_1.5 in rats (rNa_V_1.5) when applied from either side of the membrane. The blocking effect of extracellular QX-314 was reduced by substitution of DIVS6 T1755 in cardiac rNa_V_1.5 (equivalent to T1753 in hNa_v_1.5) to valine in brain rNa_V_1.2 (Qu et al., 1995). Similarly, mutation of the equivalent residue C1572 in muscle rNa_V_1.4 to threonine in cardiac rNa_V_1.5 also allowed QX-222 to block the channel from the extracellular side (Sunami et al., 2000). In addition, mutations of I1575 in DIVS6 of muscle rNa_V_1.4 or equivalent residue I1760 in brain rNa_V_1.2 (I1756 in hNa_v_1.5) to alanine (relatively small amino acid) created external access pathway for QX-222 (Sunami et al., 2001). Remarkably, these residues (T1753 and I1756 in DIVS6 in hNa_v_1.5) are part of the E2 and C1 binding sites forming the hydrophobic pathway in our simulations (Figure 4). We hypothesize that equivalent positions in other Na_V_ channels could form a hydrophobic pathway for drug access from the extracellular environment for both neutral and charged drugs. While neutral drugs may pass along the hydrophobic pathway to access the binding site within the pore lumen, charged drugs may pass along this pathway only if polar or small side chain amino acids are present in this critical region to lower the energy barrier for drug access. Results from previously published experimental data provide structural explanations for the ultra-fast blocking kinetics of extracellularly applied neutral drugs on Na_V_ channels (Hille, 1977). This hydrophobic drug access pathway in our simulations also revealed another interesting observation. Neutral lidocaine is climbing down the vertical lipid - channel interface formed by the P1-helix in DIII, P2-helix in DIV, and DIII-DIV fenestration (Supplemental Movie 2). Since neutral lidocaine is amphipathic, this could be considered to be an energetically favorable pathway. We hypothesize that other ion channels and transmembrane proteins can adopt a similar amphipathic drug access pathway at the interface between lipid and protein environments.

### Molecular dynamics simulations reveal two *neutral* lidocaines simultaneously binding within the hNa_v_1.5 channel pore lumen

Our unbiased simulations of neutral lidocaine revealed up to two lidocaine molecules binding within the channel pore lumen (Figure 5). When there is one molecule in the pore, neutral lidocaine is localized at two district binding sites NA1 and NA2. NA1 is the binding site at the center of the pore, involving Y1767 and other residues from the S6 segment of all four domains. There are limited contacts of neutral lidocaine with F1760 in the NA1. The NA2 binding site is positioned on top of F1760, near the DIII-DIV fenestration and under the P1 helix in DIII, which is similar to the most frequent and lowest interface energy pose for neutral lidocaine observed by RosettaLigand (Figure 2A). Both the amine group and the phenyl ring of lidocaine form interactions with F1760, L1462 and I1466. Two lidocaine molecules in the pore can occupy both of the NA1 and NA2 binding sites, which are sampled by a sole lidocaine molecule in the pore (Figure 5). The first neutral lidocaine in our model is positioned in a binding site formed by a region above F1760 and under the P1-helix in DIII, and fenestration region between DIII-DIV, i.e. a site equivalent to NA1 for one lidocaine in the pore. The second neutral lidocaine is positioned between F1760 and Y1767 in the central pore, resembling a single lidocaine NA1 binding site. We classify them in general as DIVS6 F1760 binding site and central pore binding site. While the lidocaine binding at F1760 is unchanged during simulations, lidocaine binding at the central pore can shift up and down, thus creating two states of binding NB1 and NB2 (Figure 5). These observations from our simulations are in agreement with experimental data showing that F1760 and Y1767 in hNa_v_1.5 play key roles in lidocaine binding (Ragsdale et al., 1996). It is also noticeable that the DIII selectivity filter region residue K1419 is part of the “DEKA” motif and plays an important role in Na_V_ channel selectivity (Hilber et al., 2005; Perez-Garcia et al., 1997). Mutations of K1419 to serine or glutamate enhance slow inactivation of Na_V_ channels (Todt et al., 1999). It is possible that while binding at the central pore can provide a simple steric blocking mechanism, lidocaine binding at F1760 and the P1-helix in DIII may directly interfere with the normally conductive state of the selectivity filter region and induce a conformational change that may promote transition to the slow inactivated state. Remarkably, cooperative binding of multiple lidocaine molecules to Na_V_ channels have been previously suggested based on dose response of inhibition with a Hill coefficient value greater than 1 (Leuwer et al., 2004). Furthermore, N-linked lidocaine dimers have been previously shown to bind to Na_V_ channels with 10–100-fold higher affinity than lidocaine monomers (Smith et al., 2006). These experimental observations agree with our MD simulation results and suggesting that lidocaine may have at least two “hot spots” for binding within the Na_V_ channel pore lumen formed between the P1 helix from domain III, F1760, and Y1767.

**Figure 5.**
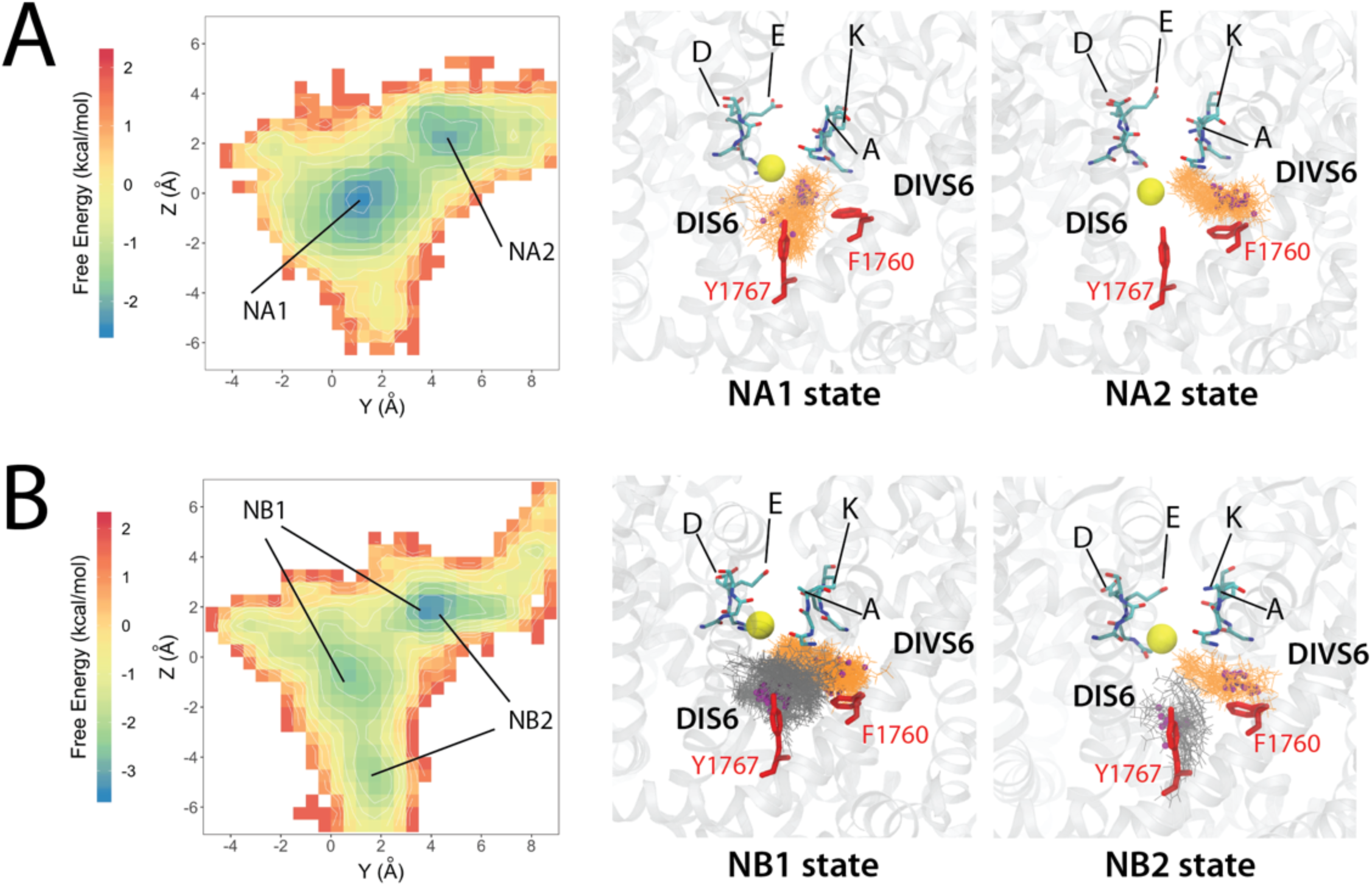
Molecular dynamics simulation of the hNa_v_1.5 channel interaction with neutral lidocaine reveal two binding poses: (A) states NA1 and NA2 for one lidocaine bound in the pore lumen; (B) NB1 and NB2 for two lidocaine molecules binding in the pore lumen at the same time. Left panels show free energy surfaces projected on the *yz* plane with binding sites identified from free energy minima and labeled. Middle and right panels show close-up transmembrane views of molecular models of charged lidocaine binding. In the close-up views lidocaine molecules (orange and dark-gray) and interacting residues on the channel (red) as well as SF “DEKA” motif (cyan for C, blue for N and red for O) are shown using stick representation. Lidocaine basic N atoms are shown as small purple spheres, and a SF bound Na^+^ atom is shown as yellow sphere.

### Molecular dynamics simulations reveal two unique “hot spots” for binding of *charged* lidocaine in the hNa_v_1.5 channel pore lumen

Unbiased MD simulations of high concentrations of charged lidocaine molecules placed in aqueous solution have shown that the drug did not pass either through the opening formed by the hydrophobic intracellular gate or through the fenestration between domains III and IV during 1 μs simulation (data not shown). Combined with results from our calculation of charged lidocaine membrane partitioning above, we suspect that those events may not be effectively sampled in a few microseconds simulation time. To further understand interactions of charged lidocaine with the hNav1.5 channel, we explored potentially unique binding poses by starting simulations with one or two charged lidocaines in the pore lumen, as described in Materials and Methods.

Simulation of one charged lidocaine revealed two highly convergent binding states lining along the vertical pore axis with the protonated amine (i.e. cationic ammonium) group of lidocaine in close proximity to the DI and DII selectivity filter region and the phenyl group of lidocaine pointing down into the lumen (see CA1 and CA2 states in Figure 6A). The CA1 state represents binding of charged lidocaine at the central pore with the protonated amine group attracted to the electron negative region below the selectivity filter. Interestingly, most of the time during the simulation, lidocaine binding in CA1 appeared to have a sodium ion binding in the selectivity filter, right above the protonated amine group. Whereas, in the absence of sodium binding in the CA2 state, charged lidocaine binds directly to the selectivity filter with the sodium binding site being taken by the protonated amine. We found that this result highly agrees with a variety of functional, structural and computational data suggesting that the selectivity filter region may form a part of local anesthetic drug binding (Bagneris et al., 2014; Buyan et al., 2018; Sunami et al., 1997; Tikhonov & Zhorov, 2017). However, compared to single neutral lidocaine binding result, we did not see the involvement of F1760 in binding of one charged lidocaine. We assume this is a result of limited sampling from 1μs unbiased simulation, although a similar result was observed in the simulation of charged lidocaine with open Na_V_Ms and closed Na_V_PaS channel using an enhanced sampling technique of replica exchange solute tempering (Buyan et al., 2018).

**Figure 6.**
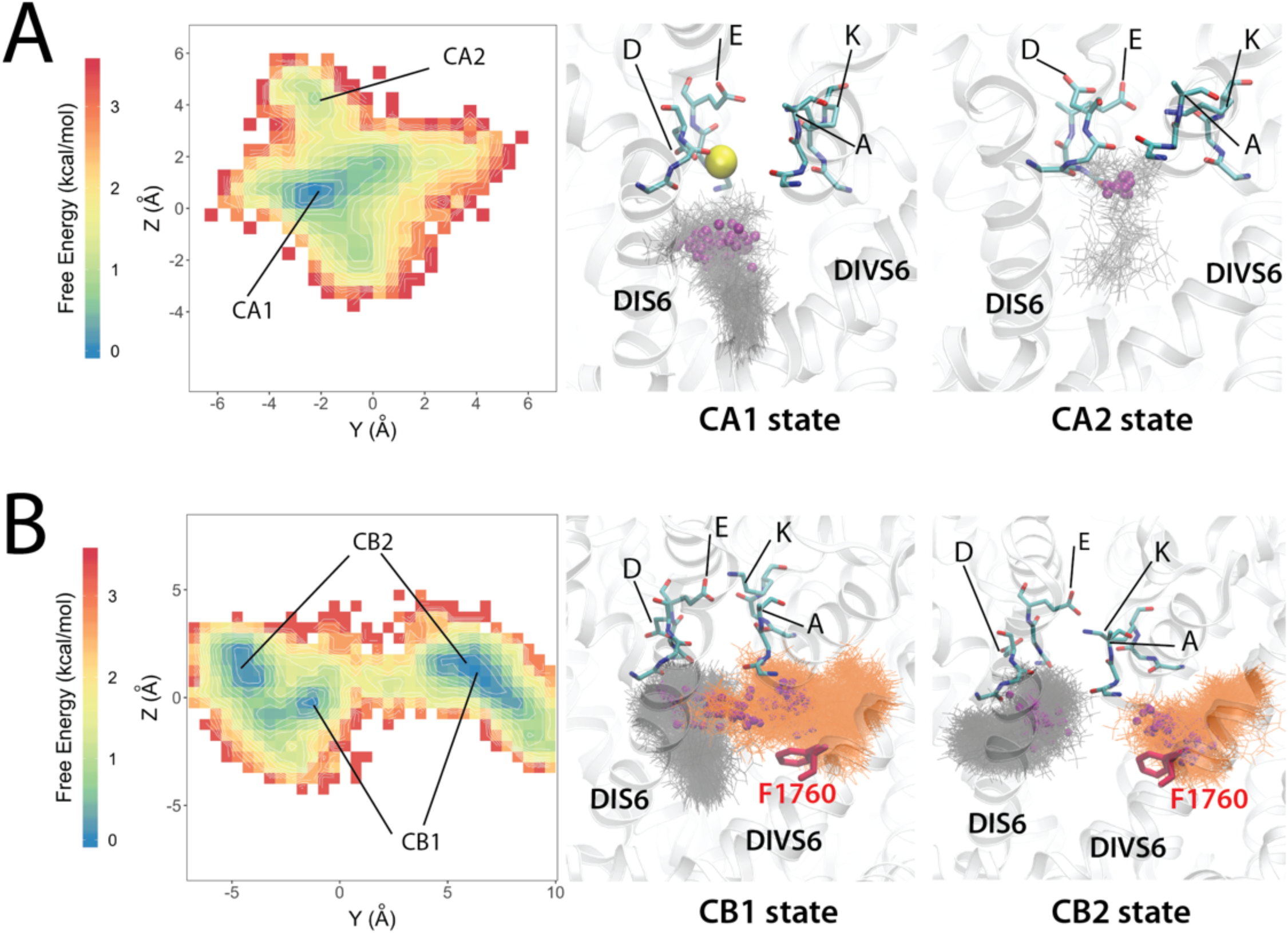
Molecular dynamics simulation of hNa_v_1.5 channel interaction with charged lidocaine reveal two binding poses: (A) states CA1 and CA2 for one lidocaine bound in the pore lumen; (B) CB1 and CB2 for two lidocaine molecules binding in the pore lumen at the same time Left panel shows free energy surface projected on the *yz* plane with binding sites identified from free energy minima and labeled. Middle and right panels show close-up transmembrane views of molecular models of charged lidocaine binding. Selectivity filter “DEKA” motif residues are shown in stick representation and colored in cyan for C, blue for N and red for O. Sodium ions are shown as spheres and colored in yellow. Lidocaine molecules are shown in stick representation and colored in gray or orange. The nitrogen atoms of the tertiary ammonium groups on charged lidocaine molecules are shown as small spheres and colored in purple. The F1760 sidechain is shown in stick representation and colored in red.

Simulation of two charged lidocaines revealed two localized binding sites, a DIVS6 F1760 binding site and a central pore binding site, similar to the case of neutral lidocaine. While lidocaine binding at the F1760 site is relatively stable, binding to the central pore can be shifted creating two highly convergent states, CB1 and CB2 (Figure 6). The first highly converged state (CB1) has one charged lidocaine lining along the vertical pore axis with the protonated amine group in close proximity to the DI and DII selectivity filter region and the phenyl group pointing down into the lumen (see CB1 state in Figure 6B), the same orientation as for one lidocaine molecule (CA1 state in Fig. 6A). Another charged lidocaine at the DIVS6 F1760 site has the protonated amine group forming cation-π interactions with F1760 and the phenyl group pointing into the fenestration region between DIII and DIV (see CB1 state in Figure 6B). Notably, the cation-π interaction is dominant during the simulation. We rarely observed π-π stacking interactions between the phenyl ring of charged lidocaine and F1760. This agrees with experimental data suggested that interactions between charged lidocaine and F1760 are cation-π interactions, not π-π interactions (Ahern et al., 2008). The second highly converged binding state (CB2) has the central pore localized charged lidocaine oriented mostly along the horizontal membrane plane (not the vertical transmembrane axis as in CB1) with the protonated amine group also in close proximity to the DI and DII selectivity filter region. However, the phenyl group is pointing into the fenestration region between DI and DII (see CB2 state in Figure 6B). The other charged lidocaine at the DIVS6 F1760 site forms an interaction with F1760 in a similar manner to that in the CB1 state.

It is interesting to note that F1760 has been shown to be a key determinant for the use-dependent block while Y1767 only has a modest effects (Ragsdale et al., 1994). In addition, mutation of W1531 to Cys in Nav1.4 (W1713 in our hNav1.5) in the DIV-P2 region was shown to abolish use-dependence of mexiletine and QX-222, without destabilizing fast inactivation or altering drug access (Tsang et al., 2005). In our model, W1713 is part of the binding site E2 for the neutral lidocaine pathway (Figure 4) and is the ceiling of the DIII-DIV fenestration, right above F1760. The best RosettaLigand docking models, MD simulations of both neutral and charged lidocaine identified the DIVS6 F1760 site as a common binding site. Together, these results encourage us to propose the binding site at DIVS6 F1760, near the DIII-DIV fenestration as the high affinity use-dependent binding site. Whereas, other binding sites at the selectivity filter region (for charged lidocaine) and at central pore near Y1767 (for neutral lidocaine) can be considered based on our simulations as low affinity binding sites. Tonic block was not the focus of this study and may require investigation of interactions with the channels in a resting state. However, because of the modest effect of F1760 and W1713 on tonic block (Ragsdale et al., 1994, 1996; Tsang et al., 2005), it may not be surprising if the tonic block binding site is similar to one of the low affinity binding sites we observed here for the interaction of lidocaine with a putatively inactivated state channel.

### Lidocaine binding to hNa_v_1.5 attenuates sodium binding in the selectivity filter

MD simulations of the hNa_v_1.5 channel in the absence or presence of 1 or 2 neutral or charged lidocaine molecules suggest that binding of lidocaine to its receptor site(s) within the pore lumen reduces Na^+^ ion binding within the selectivity filter region (Figure 7). Free energy surfaces for a Na^+^ ion within the hNa_v_1.5 selectivity filter reveal 3 major Na^+^ binding sites within this region (see sites S1, S3, and S3 in Figure 7) and 1 additional site within the pore lumen (see site S0 in Figure 7). Site S1 is located just below the selectivity filter region and formed by the carbonyl groups of T370 and Q371 (in DI) and C896 and G897 (in DII). Site S2 is formed by the carboxylate groups of D372 (in DI) and E898 (in DII) - residues in the classical “DEKA” selectivity filter motif in Na_V_ channels. Site S3 is formed by the carboxylate groups of E375 (in DI), E901 (in DII), D1423 (in DIII), and D1714 (in DIV). In the absence of lidocaine, all 3 Na^+^ binding sites are well-defined (Figure 7A and B). When 1 or 2 neutral lidocaine molecules are present in the pore lumen, the Na^+^ binding site S1 diffuses further into the pore lumen region, while sites S2 and S3 within the selectivity filter region are occupied less frequently (Figure 7A and B). When 1 or 2 charged lidocaine molecules are present in the pore lumen, we observe a dramatic reduction in Na^+^ binding at the pore lumen site S0 and within the selectivity filter region in all 3 sites, especially at sites S2 and S3 (Figure 7A and B). This disruption of continuous ion density in those cases (see Figure 7A) may impair ion conduction through the selectivity filter.

**Figure 7.**
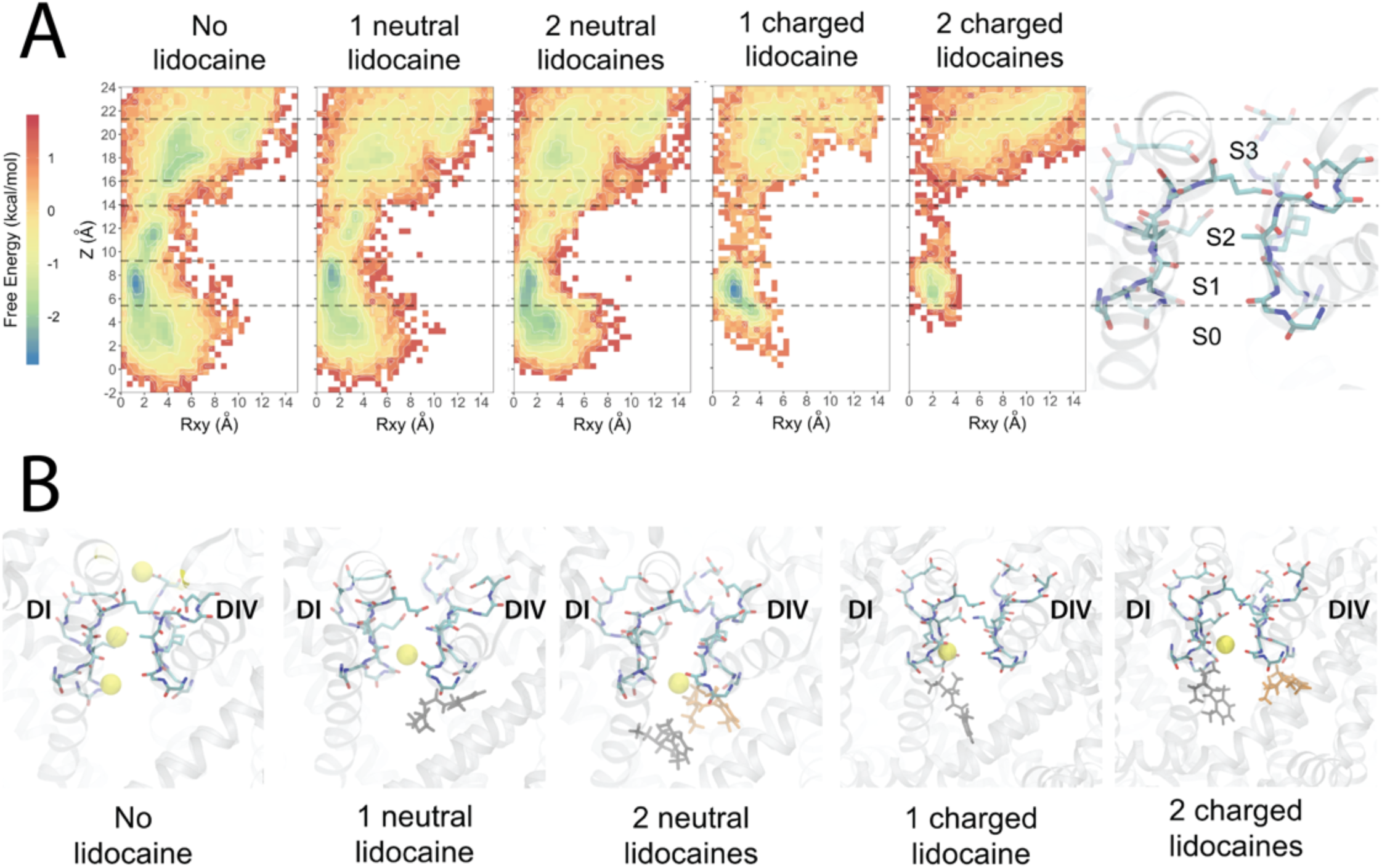
Molecular dynamics simulations reveal the free energy surfaces and binding sites for sodium ion within hNa_v_1.5 pore. (A) Transmembrane view projection of the free energy surface for sodium ion without lidocaine and in the presence of 1 or 2 neutral or charged lidocaine molecules. Specific Na^+^ binding sites are labeled S0, S1, S2, and S3 in the molecular representation of the channel SF on the right panel. (B) Representative transmembrane views of sodium ion binding sites within the selectivity filter region of the channel observed without lidocaine and in the presence of 1 or 2 neutral or charged lidocaine molecules. Sodium ions are shown as yellow spheres. The selectivity filter region residues are shown in stick representation and labeled.

The positioning of neutral or charged lidocaine molecules under the selectivity filter region in the MD simulations is notable with respect to experimental data that have identified specific mutations in the selectivity filter region that significantly affect the slow inactivation of Na_V_ channels (Balser et al., 1996; Kambouris et al., 1998; Ong et al., 2000; Todt et al., 1999). Interestingly, decreasing extracellular [Na^+^] potentiates use-dependent block by lidocaine (Chen et al., 2000). Lidocaine binding under the selectivity filter region may induce conformational changes in the selectivity filter that may enhance slow inactivation (Chen et al., 2000; Fukuda et al., 2005). However, raising extracellular [Na^+^] inhibits native slow inactivation of Na_V_ channels (Chen et al., 2000).

### Conclusions

Our structural modeling and simulation of antiarrhythmic and local anesthetic drugs interacting with the human Na_V_1.5 channel revealed the following key observations: (1) The region above F1760 in the DIVS6 segment forms a “hot spot” for drug binding and extends from the fenestration region between the DIIIS6 and DIVS6 segments to the hydrophobic pockets under the selectivity filter regions in DIII and DIV; (2) The amine/ammonium group of lidocaine, etidocaine, and QX-314 is positioned above and near F1760 (Figure 2). The phenyl ring of lidocaine, etidocaine, and QX-314 is observed in multiple different orientations near F1760 (Figure 2); (3) Flecainide and ranolazine bind to a larger protein surface area that spans from the fenestration region between the DIII and DIV to the ion conduction pathway under the selectivity filter region; (4) Lidocaine enters the hNa_v_1.5 pore via the hydrophilic pathway through the intracellular gate and via a hydrophobic pathway through a fenestration between DIIIS6 and DIVS6 (Figure 4); (5) up to two lidocaine molecules can simultaneously bind within the hNa_v_1.5 pore lumen (Figures 5 and 6); (6) bound lidocaine molecules can interfere with the ion occupancy in the hNa_v_1.5 SF (Figure 7).

Our results provide crucial atomic scale mechanistic insights into protein - drug interactions, necessary for the rational design of novel modulators of the cardiac Na_V_ channel to be used for the treatment of cardiac arrhythmias. The fundamental novelty of bringing together Rosetta molecular modeling and MD simulations to study drug - channel interactions has the potential to enable automated virtual drug screening in the future. Critically, this approach can be applied to any ion channel, which might be used to predict inDIVidual patient responses to drug therapy based which specific ion channel mutations they have. For instance, we can predict how a single mutation in ion channel encoding gene would affect protein - drug binding and how an effect of such alteration propagates from a protein to a single cell and the cardiac rhythm of the whole organ. This work sets the stage for expansion to novel linkages by connecting mature experimental structural and functional approaches to emerging modeling approaches at the atomic and organ scales. There is potential for future simulations to be carried out to predict how functional properties of drugs can be perturbed in an emergent multiscale modeling system, and these predictions may ultimately be used to inform structural models to screen drug analogs that confer the requisite functional properties predicted critical for therapy.

In particular, this study represents the first critical step for elucidating structural determinants of drug cardiac safety profiles at atomic resolution. We have observed differences in Na_V_1.5 binding profiles for cardiac safe lidocaine versus flecainide, a drug with a known proclivity for deadly arrhythmia. Our previous multi-scale modeling and experimental study suggested that such molecular scale differences can propagate and emerge at the tissue and organ levels as notable pro-arrhythmia markers (Moreno et al., 2011). We have also performed multi-microsecond molecular dynamic simulations to explore drug - channel binding pathways for charged and neutral forms of lidocaine, which provided a molecular picture consistent with previous experimental observations. Future work will extend these studies to flecainide and other Na_V_1.5 channel binders with different pro-arrhythmia proclivities.

## Materials and Methods

### Rosetta modeling of the hNa_v_1.5 channel

We used the Rosetta structural modeling software (Alford et al., 2017; Bender et al., 2016; Rohl et al., 2004) and the cryoEM structure of the Na_V_1.4-beta1 complex from the electric eel (eeNa_V_1.4) (PDB ID: 5XSY) as a template to predict the structure of the human Na_v_1.5 (hNa_v_1.5) channel. At first, the structure of eeNa_V_1.4 without the beta1 subunit was passed through the Cryo-EM refinement protocol in Rosetta (DiMaio et al., 2015). The lowest scoring density-refitted eeNa_V_1.4 model and electron density were then used in combination in RosettaCM (Song et al., 2013) to model the hNa_v_1.5 channel. We generated 5,000 structural models of hNa_v_1.5 and selected the top 500 lowest-scoring models for clustering analysis as described previously (Bonneau et al., 2002). Models from top clusters were visually inspected to select the final model for the docking study.

### RosettaLigand modeling of hNa_v_1.5 channel interaction with antiarrhythmic and local anesthetic drugs

OpenEye OMEGA (OpenEye Scientific Software) (Hawkins & Nicholls, 2012; Hawkins et al., 2010) was used to generate conformers for antiarrhythmic and local anesthetic drugs. To uniformly and efficiently sample the pore region of hNa_v_1.5, drugs were placed at 5 different initial locations: at the center of the cavity and at 4 fenestration sites. We incorporated an initial random perturbation with a translation distance less than 10 Å before the docking run to add another layer of randomization. Sampling radius was set to 10 Å. The details of the RosettaLigand docking algorithm have been described previously (Bender et al., 2016; Combs et al., 2013; Davis & Baker, 2009; DeLuca et al., 2015; Meiler & Baker, 2006). A total of 200,000 docking models were generated for each drug. The top 10,000 models were selected based on the total score of protein-ligand complex and then ranked by ligand binding energy represented by Rosetta interface delta_X energy term. The top 50 ligand binding energy models were visually analyzed using UCSF Chimera (Pettersen et al., 2004) and the most frequently sampled ensembles of poses are shown in Figures 2 and 3, with several representative poses demonstrated in Figure 2 and 3 Figure Supplements.

### Drug forcefield parameterization

We obtained the molecular structure of lidocaine from the ZINC database (accession number 20237), (Irwin & Shoichet, 2005), and used the CGENFF program, version 1.0 (Vanommeslaeghe & MacKerell; Vanommeslaeghe et al.) to generate initial guesses for partial atomic charges, bond lengths, bond angles, and dihedral angles.

The initial topology and parameters for charged and neutral forms lidocaine were subsequently validated and optimized using QM target data following the suggested CGENFF force field methodology (Vanommeslaeghe et al.). High-quality parameters not already present in CGENFF are assigned from existing parameters based on chemical analogy, and our optimizations focused on parameters with poor chemical analogy corresponding to a high penalty score (Vanommeslaeghe et al.). The Force Field Toolkit plugin (ffTK) (Mayne et al., 2013) for the Visual Molecular Dynamics program (VMD) (Humphrey et al., 1996) was used to generate files for quantum mechanical (QM) reference calculations and to perform parameter optimizations. QM target data for parameter optimization were obtained utilizing Møller-Plesset (MP2) and Hartree-Fock (HF) electronic structure methods and the 6–31(d) basis set using the Gaussian 09 program (Frisch et al., 2009).

MP2/6-31G(d) molecular dipole magnitude and orientation as well as scaled HF/6-31G(d) interaction energies with water were used for the optimization of partial atomic charges compatible with the CHARMM atomistic force fields (Mackerell). Internal bond and angle parameters were validated by comparison to MP2/6-31G(d) optimized geometries and scaled vibrational frequencies, and differences within 0.01 Å and 1° between QM and MM equilibrium bond and angle values were sought. Finally, the dihedral angle parameters were optimized to reproduce MP2/6-31G(d) potential energy scans for rotation around a particular bond.

Optimized charges (**Table S1**) are in good agreement with QM target dipole values. The optimized MM dipole moments are overestimated in magnitude from QM MP2/6-31G(d) dipole moments by 17% for neutral lidocaine and 16% for charged lidocaine (close to a 20% acceptable lower-end threshold, suggested for the CGENFF force field), and the MM dipole direction differed by ~1° from the QM computed direction for both charged and neutral lidocaine. The water interaction distances were all within 0.4 Å of QM target values (see **Tables S2 and S3)**. The MM dipole moment for charged lidocaine (11.68 Debye) is almost three times higher than for neutral lidocaine (3.93 Debye), which agrees with respective computed QM values. Water interaction energies were also in good agreement with QM values, with root mean squared errors (RMSE) of 0.95 kcal/mol for neutral lidocaine, and 1.41 kcal/mol for charged lidocaine, respectively (**Table S4**). For neutral lidocaine, there was a high penalty score for the C2-N1-C3 bond angle, and optimization yielded a difference of 0.16° between MM and QM values. For charged lidocaine there were no high penalties for internal bond and angle parameters from the CGENFF. For neutral lidocaine, there were four high-penalty dihedral angles, and for charged lidocaine there were two high-penalty dihedral angles from the CGENFF. Dihedral optimizations resulted in great improvement over CGENFF initial guesses (illustrated in **Figure 4 - figure supplement 1 and 2**), with optimized torsional energy minima within ~2 kcal/mol of QM values. For comparison, raw CGENFF dihedral parameters with high penalties yielded QM free energy minima differences sometimes as high ~5kcal/mol.

Final topology and parameters for neutral and charged lidocaine are provided in the Appendix.

### Drug-membrane partitioning

Partitioning of charged and neutral lidocaine into a lipid membrane was assessed using the NAMD (Phillips et al., 2005) program. Initial system setup scripts were generated with the CHARMM-GUI web toolkit (Sunhwan Jo et al., 2008) and were modified to build the hydrated drug-membrane systems, which consisted of 128 1-palmitoyl-2-oleoylphosphatidylcholine (POPC) lipids, ~7000 water molecules, 21 or 22 K^+^ and 22 Cl^−^ ions to ensure 0.15 M electrolyte concentration and overall electrical neutrality, and one drug molecule, totaling ~38,250 atoms. CHARMM36 lipid force field (Klauda et al., 2010), TIP3P water model (Jorgensen et al., 1983), standard CHARMM ion parameters (Beglov & Roux, 1994) and CGENFF (Vanommeslaeghe et al., 2010b) compatible drug parameters developed in this work were used throughout all simulations.

For partitioning calculations of each drug we used the umbrella sampling (US) method (Torrie & Valleau, 1977) with 81 independent simulation windows, placing the center of mass (COM) of a randomly oriented drug molecule in 1 Å intervals from -40 Å to 40 Å with respect to COM of the membrane. The COM of the drug was restrained along the *z* axis with a force constant of 2.5 kcal/mol/Å^2^, and an additional 5 kcal/mol/Å^2^ cylindrical restraint was applied in order to prevent the drift of the molecule in the *xy* plane. Each NAMD US simulation of charged and neutral lidocaine was carried out in a *NPT* ensemble with 1 atm pressure maintained by Langevin piston barostat (Feller et al., 1995), and 310K, controlled by Nosé-Hoover thermostat (Hoover, 1985; Nosé, 1984). Tetragonal cells with periodic boundary conditions (PBC) were used in all the simulations, and the SHAKE algorithm (Ryckaert et al., 1977) was employed to fix the bonds to all hydrogen atoms, allowing for the use of a 2 fs time step. Electrostatic interactions were computed via Particle Mesh Ewald (Darden et al., 1993), with a mesh grid of 1 Å. Potential of mean force (PMF) profiles were computed using the weighted histogram analysis method (WHAM) (Kumar et al., 1992). Umbrella sampling simulations for charged and neutral lidocaine were run for 15 ns per window.

Drug-water partition coefficients were calculated as was done previously (Vorobyov et al., 2012):

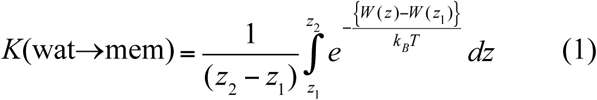

where W(*z*) is the PMF, *z*_1_ and *z*_2_ are points in aqueous solution on opposite sides of the membrane, k_B_ is Boltzmann constant, and *T* is the absolute temperature.

Error bars were estimated from PMFs by propagation of uncertainties.

The distribution coefficient, *D*, was computed as

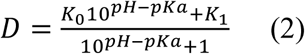

Where *K_0_* is the partition coefficient of a neutral drug form, and *K_1_* is the partition coefficient of a charged (protonated) drug form, both computed via Equation 1.

To compute drug translocation rates across membrane we used Kramer’s transition rate approximation as was done previously (Allen et al., 2003; Crouzy et al., 1994). For charged lidocaine local diffusion near the membrane center was computed to be *D*(*z*_barrier_)=0.0047 Å^2^/ps, and the curvatures of the PMF well and the PMF peak were 0.0508 and -0.207, respectively. For neutral lidocaine D(*z*_barrier_)=0.0089 Å^2^/ps, and the curvatures of the PMF well and the PMF peak were 0.0312 and -0.0784, respectively.

### Molecular dynamics simulations of hNa_v_1.5 channel interaction with lidocaine

The hNa_V_1.5 model was embedded in a bilayer of POPC with explicit TIP3P water molecules and 150 mM (with lidocaine) or 500 mM (without lidocaine) of NaCl using CHARMM-GUI (S. Jo et al., 2008). For lidocaine containing simulations we used physiological NaCl concentration, but we used larger salt concentration in the drug-free runs to facilitate Na^+^ conductance. For all these simulations, we also used CHARMM36 lipid (S. Lee et al., 2014) and protein (Huang & MacKerell, 2013) force fields, and CHARMM generalized force field (CGENFF) compatible parameters for lidocaine as described above. Initial system equilibrations were performed using NAMD on a local GPU cluster. After 10,000 steps of steepest descent minimization, MD simulations started with a timestep of 1 fs with harmonic restraints initially applied to protein heavy atoms and some lipid tail dihedral angles. These restraints were slowly released over 2 ns. Harmonic restraints (0.1 kcal/mol/Å^2^) were then applied only to protein Cα atoms, and the systems were equilibrated further for 50 ns with a timestep of 2 fs. In order to use a 2 fs timestep, all bonds to H atoms were constrained using the SHAKE algorithm. All simulations were performed at constant pressure (1 atm) with constant ratio of *x* and *y* dimensions in order to maintain the correct area per lipid, and constant temperature of 303.15 K (chosen to avoid the gel phase transition of POPC lipids). Electrostatic interactions were computed using Particle Mesh Ewald (PME). Non-bonded pair lists were updated every 10 steps with a list cutoff distance of 16 Å and a real space cutoff of 12 Å with energy switching starting at 10 Å.

Equilibrated systems were simulated on the Anton 2 supercomputer using Anton 2 software (Shaw et al., 2014) version 1.31.0 in the *NPT* ensemble at 303.15 K. A 2 fs timestep was used with non-bonded long-range interactions computed every 6 fs using the RESPA multiple time step algorithm. The multi-integrator (multigrator) algorithm was used for temperature and semi-isotropic pressure coupling. Long-range electrostatic interactions were handled by u-series algorithm (Shaw et al., 2014). A long-range Lennard-Jones (LJ) correction (beyond cutoff) was not used as was suggested for CHARMM36 lipid force field. For the simulation of hNa_V_1.5 without drugs, an electric field was applied downwardly in the *z* direction to mimic membrane potential of 250 mV (positive inside).

For the neutral lidocaine simulations, two different systems were created with initial neutral lidocaine aqueous concentration at 75mM and 150mM. Each system was simulated for 7 μs on Anton2.

For the charged lidocaine simulations, systems of 1 and 2 charged lidocaine were created by initially placing 1 and 2 charged lidocaine molecules in the cavity of the hNav1.5 model. Each system was simulated for 1 μs on Anton2.

## Analysis

### Drug binding in the channel

3D density maps of the drug center of mass for the neutral lidocaine and position of the amino group for the charged one from Na_V_1.5 - drug flooding MD simulations were used to compute free energy profiles using equation *W*(*r_i_*) *= -k_B_Tln*[*ρ*(*r_i_*)] *+ C* where *ρ*(*r_i_*) is the unbiased probability distribution as a function of reaction coordinates *r_i_*·, and *C* is a constant. The maps were offset to get an average free energy of 0 kcal/mol in bulk water for neutral lidocaine or for the binding site in the pore for the charged lidocaine. 2D projections of these free energy maps on the *Z* (transmembrane) and *Y* (lateral) axes are shown in Figures 4, 5 and 6. Origin is selected as the center of mass of the protein.

### Sodium binding in the selectivity filter (Figure 7)

*xy*-radial position ≤ 15Å, and z-axial position between -15 and +15 Å were used to define the pore region for ion occupation. x, y and z are defined relative to the center of mass (COM) of the backbone of the selectivity filter. Free energy surfaces were calculated from unbiased simulation as *W*(*r_i_*) *= -k_B_Tln*[*ρ*(*r_i_*)] *+ C* where *ρ*(*r_i_*) is the unbiased probability distribution as a function of reaction coordinates *r_i_*, and C is a constant. Origin is selected as the center of mass of the protein.

## Acknowledgements

We would like to thank Drs. Jon Sack, Toby Allen, Heike Wulff, Kazuharu Furutani, and members of Clancy, Yarov-Yarovoy and Sack laboratories for helpful discussions. We thank Dr. Nieng Yan (Princeton University) for sharing coordinates of electric eel and human Na_V_1.4 channel structures. Anton 2 computer time was provided by the Pittsburgh Supercomputing Center (PSC) through Grant R01GM116961 from the National Institutes of Health. The Anton 2 machine (Shaw et al., 2014) at PSC was generously made available by D.E. Shaw Research. This research was supported by National Heart, Lung, and Blood Institute Grant U01HL126273, R01HL128537, R01HL128170 to CEC and American Heart Association Predoctoral Fellowship 16PRE27260295 to KRD.

---

**Figure 1 - figure supplement 1.**
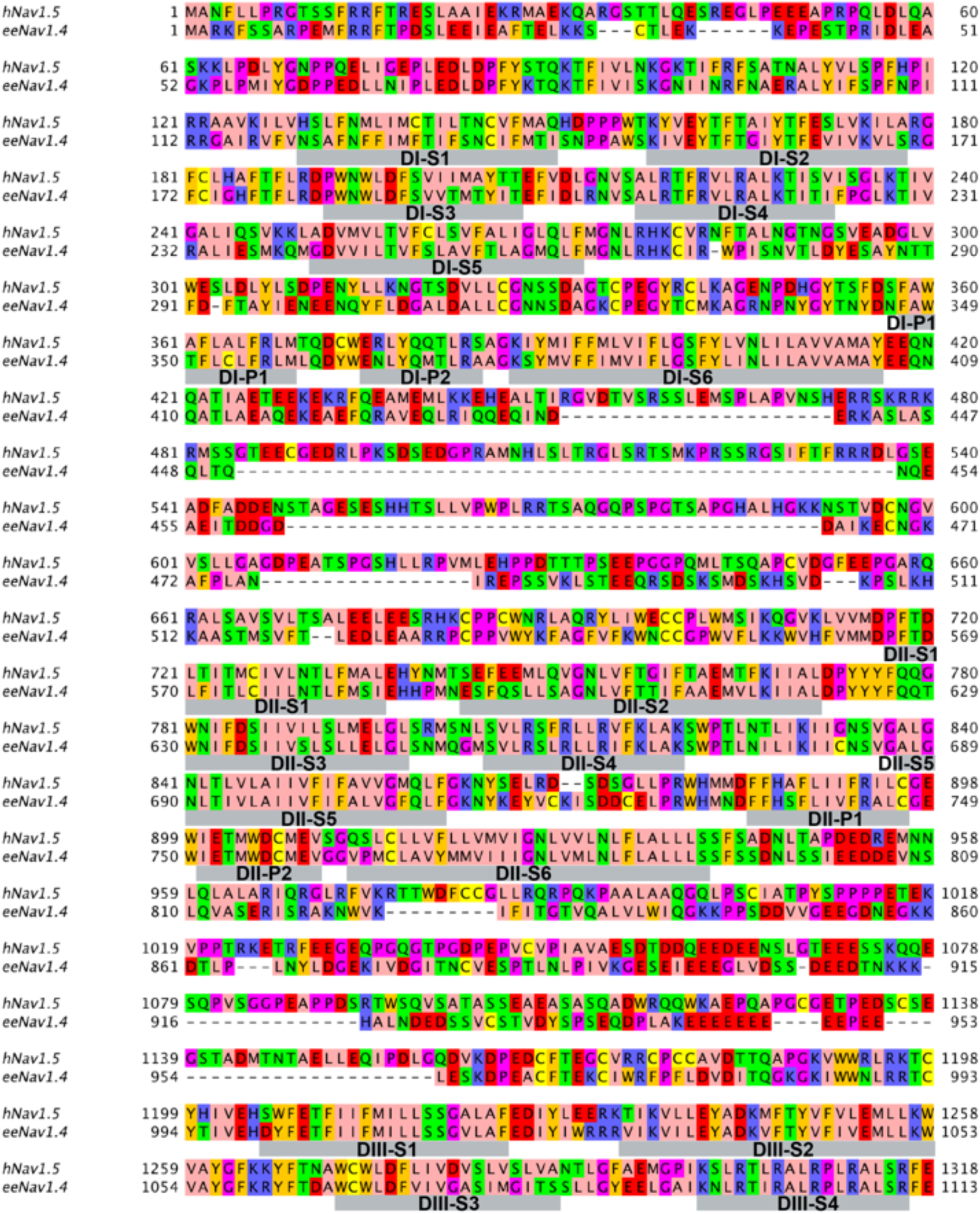
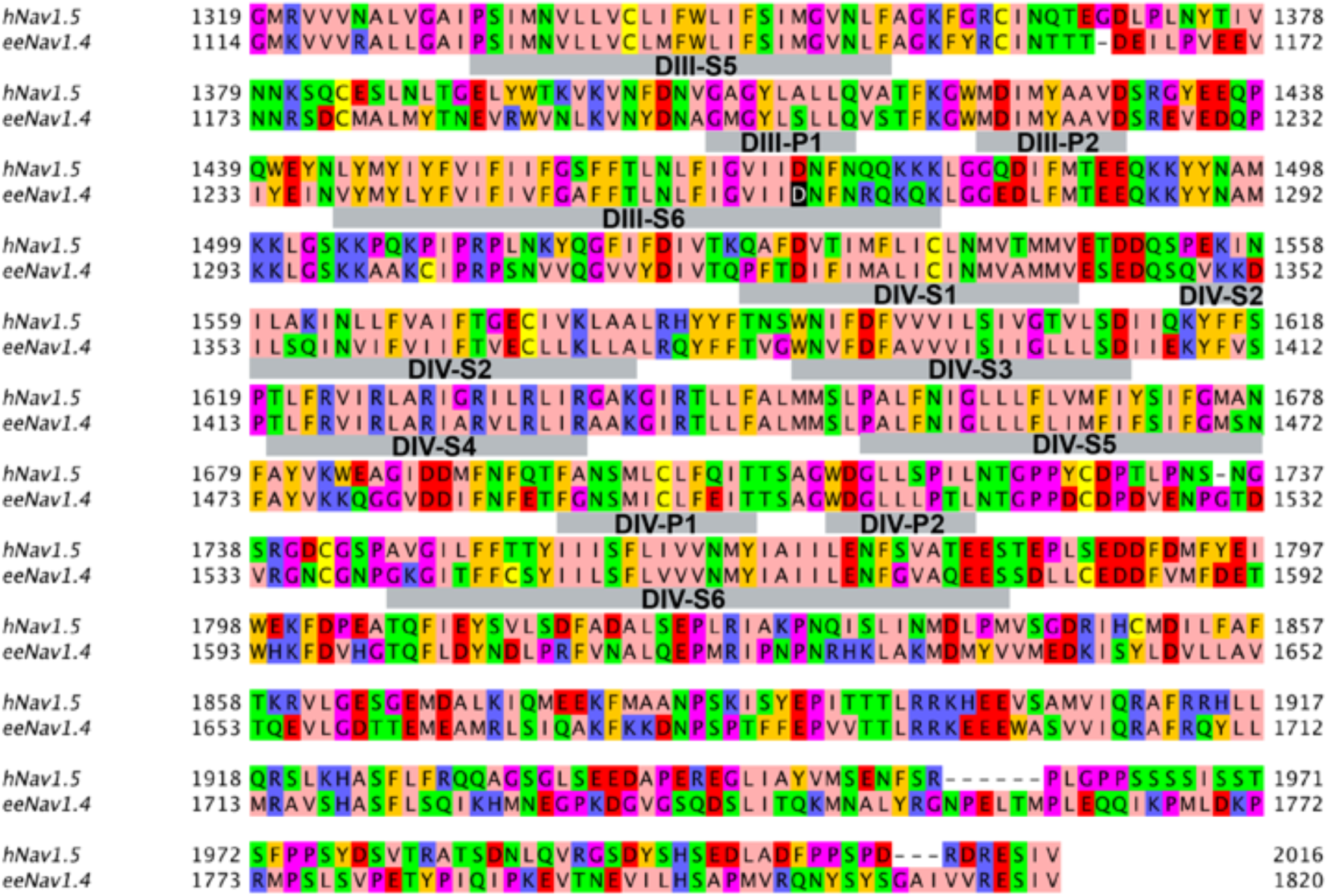
Sequence alignment between hNav1.5 and eeNav1.4. Transmembrane segments S1-S6 and P1 and P2 helix regions in each domain are underlined by gray bars and labeled. Amino acids were colored with Jalview program using the Zappo color scheme, where hydrophobic residues (I, L, V, A, and M) are colored pink, aromatic residues (F, W, and Y) are colored orange, positively charged residues (K, R, and H) are colored blue, negatively charged residues (D and E) are colored red, hydrophilic residues (S, T, N, and Q) are colored green, P and G colored magenta, and C is colored yellow.

**Figure 1 - figure supplement 2.**
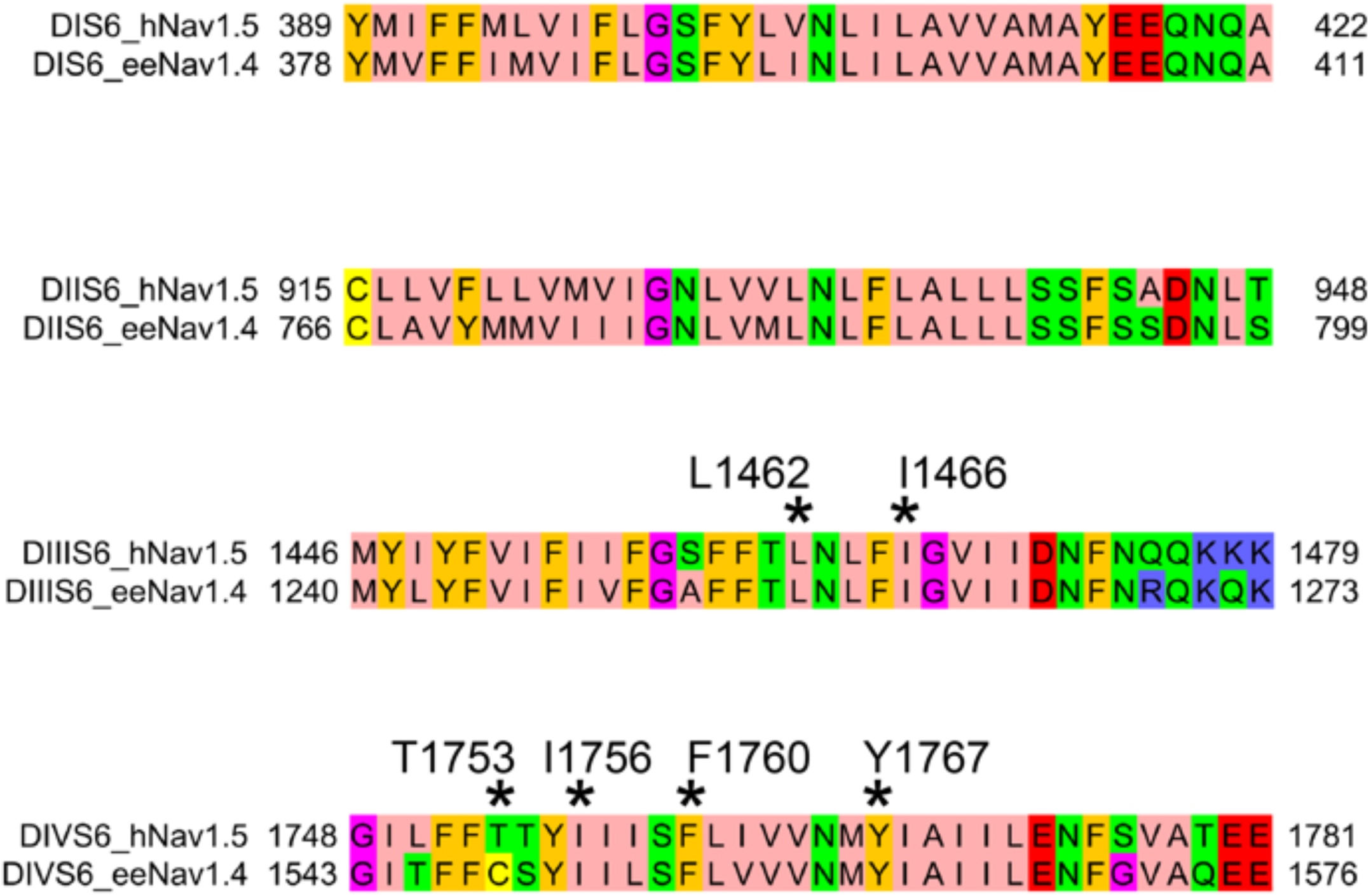
Sequence alignment between hNav1.5 and eeNav1.4 transmembrane segments S6. Specific hNav1.5 residues discussed in the main text are marked by asterisk and labeled. Amino acids were colored as in Figure 1 - figure supplement 1.

**Figure 1 - figure supplement 3.**
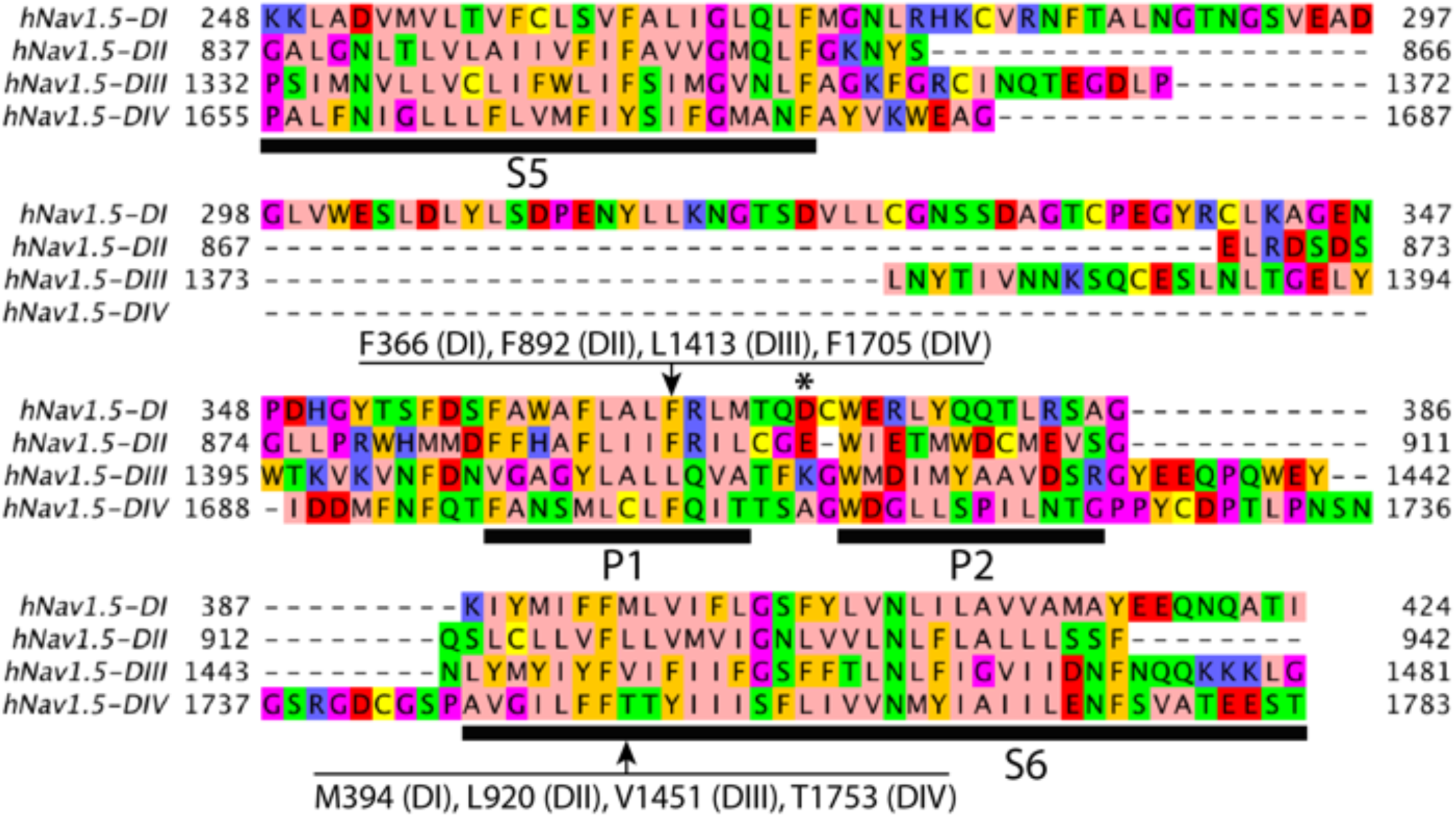
Sequence alignment between four domains of hNav1 .5 segments S5, P1-helix, P2-helix, and S6. Specific hNav1.5 residues discussed in the main text are marked by arrows and labeled. Transmembrane segments S5 and S6 and P1 and P2 helix regions in each domain are underlined by black bars and labeled. Amino acids were colored as in Figure 1 - figure supplement 1.

**Figure 1 - figure supplement 4.**
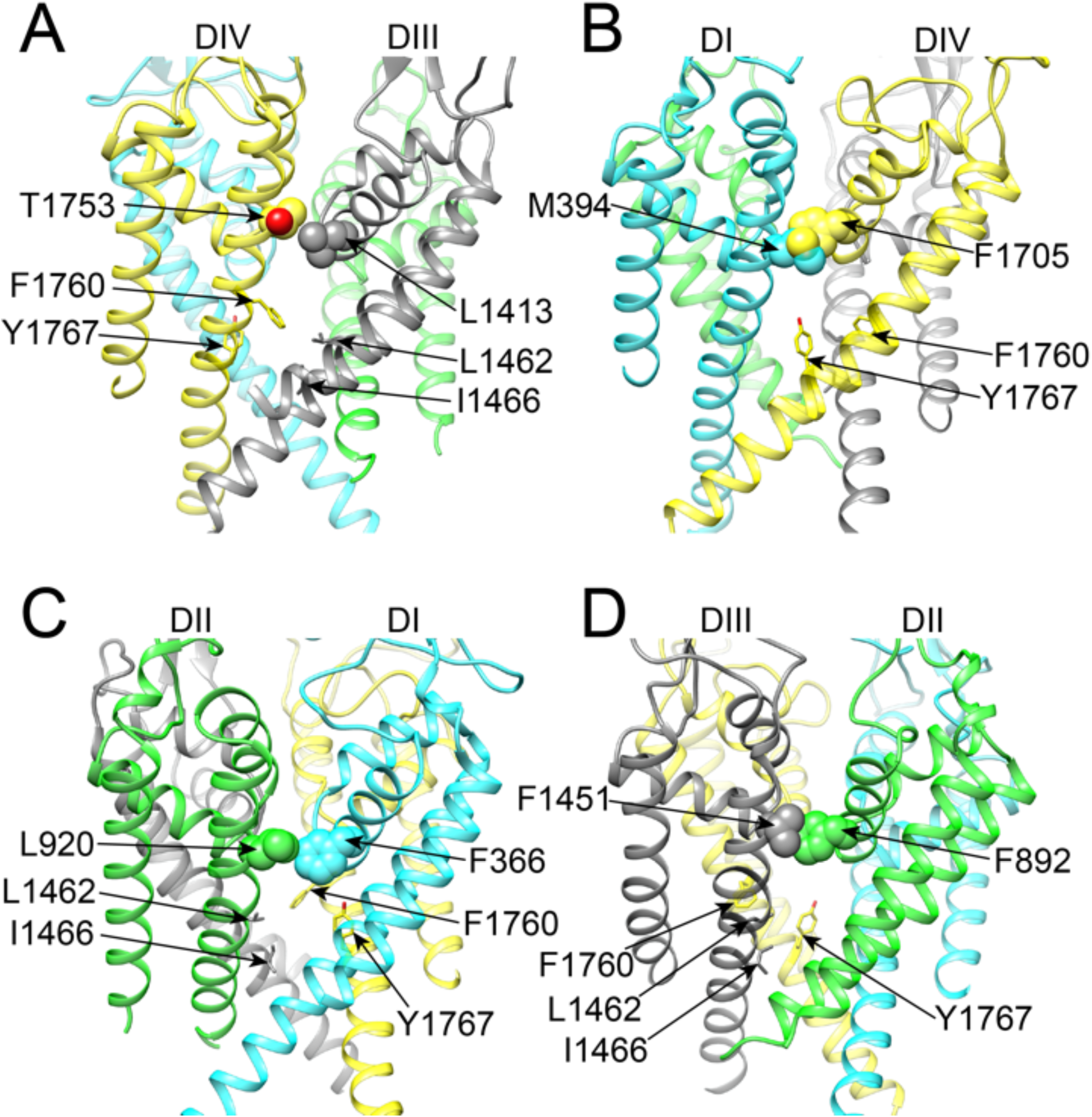
Transmembrane views of all four hNav1.5 fenestrations. (A) DIII and DIV fenestration. (B) DI and DIV fenestration. (C) DI and DII fenestration. (D) DII and DIII fenestration. Side chains of fenestration-forming residues are shown in space-filling or stick representations, labeled, and colored using corresponding domain colors, with O atom shown in red.

**Figure 2 - figure supplement 1.**
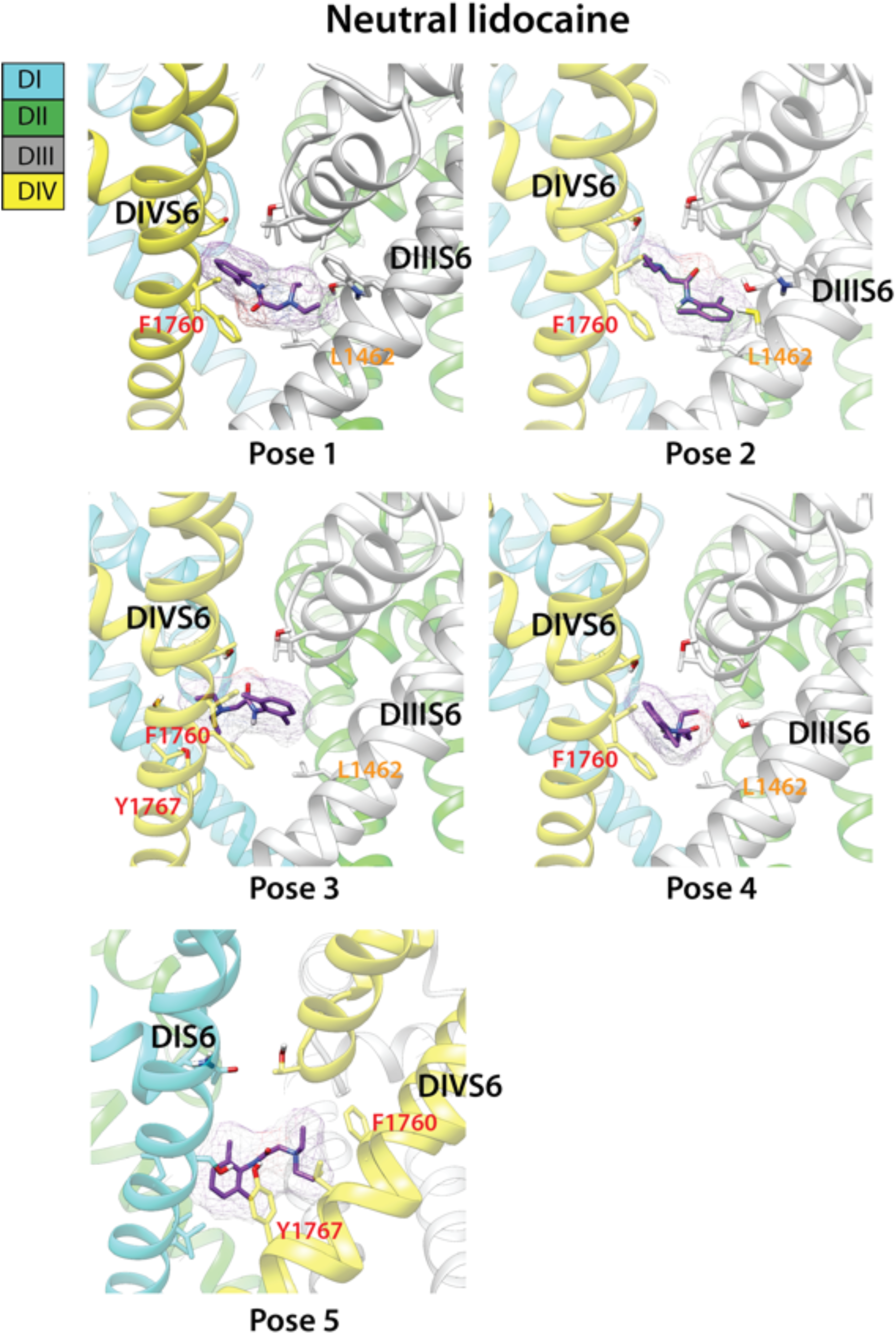
Top binding poses of neutral lidocaine interaction with Rosetta model of hNav1.5 channel. Domain I is colored in blue, domain II is colored in green, domain III is colored gray, and domain IV is colored yellow. hNav1.5 residues forming interactions with lidocaine are shown in stick representation and labeled. Lidocaine is shown in stick and surface representation and colored purple.

**Figure 2 - figure supplement 2.**
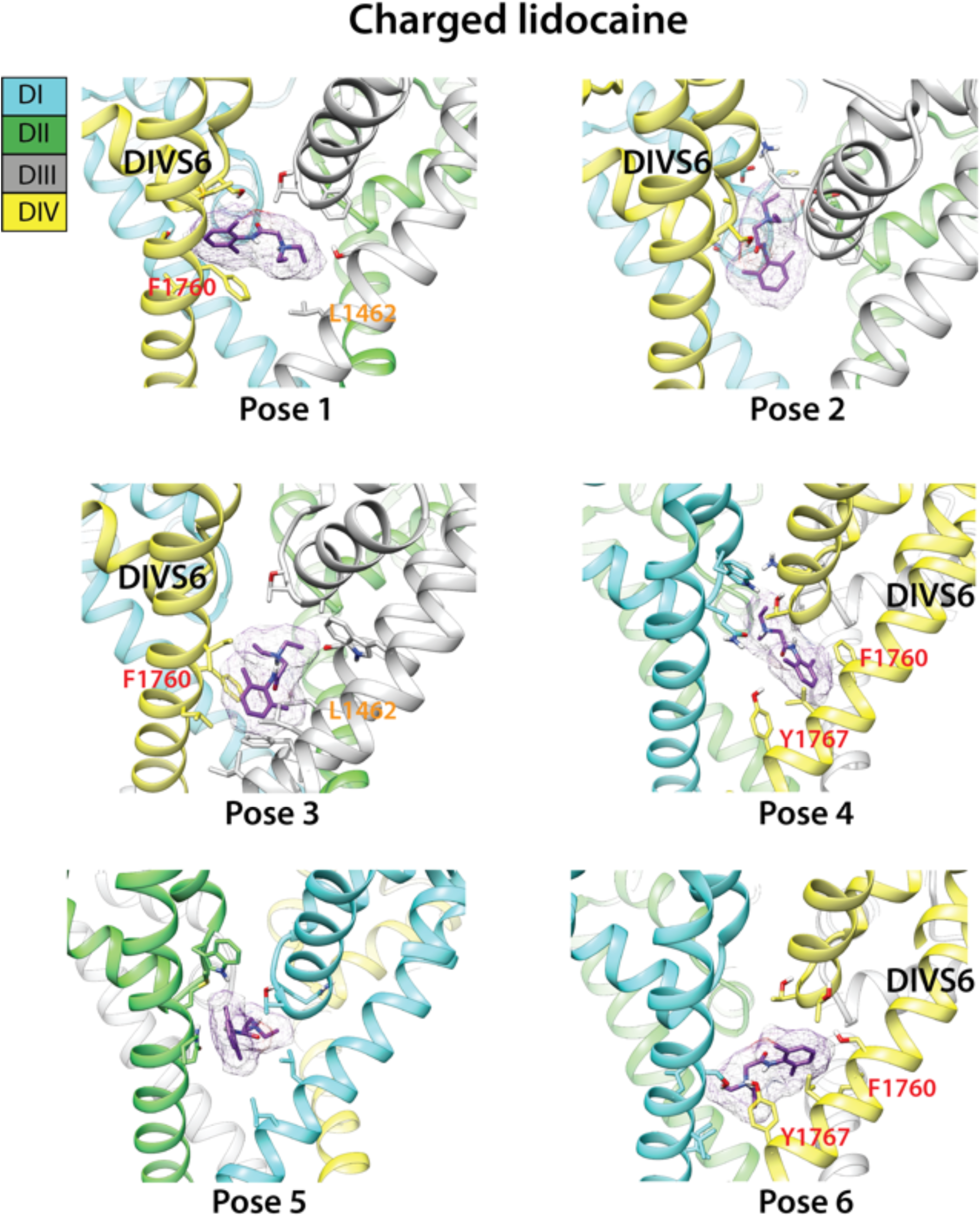
Top binding poses of charged lidocaine interaction with Rosetta model of hNav1.5 channel. Domain I is colored in blue, domain II is colored in green, domain III is colored gray, and domain IV is colored yellow. hNav1.5 residues forming interactions with lidocaine are shown in stick representation and labeled. Lidocaine is shown in stick and surface representation and colored purple.

**Figure 2 - figure supplement 3.**
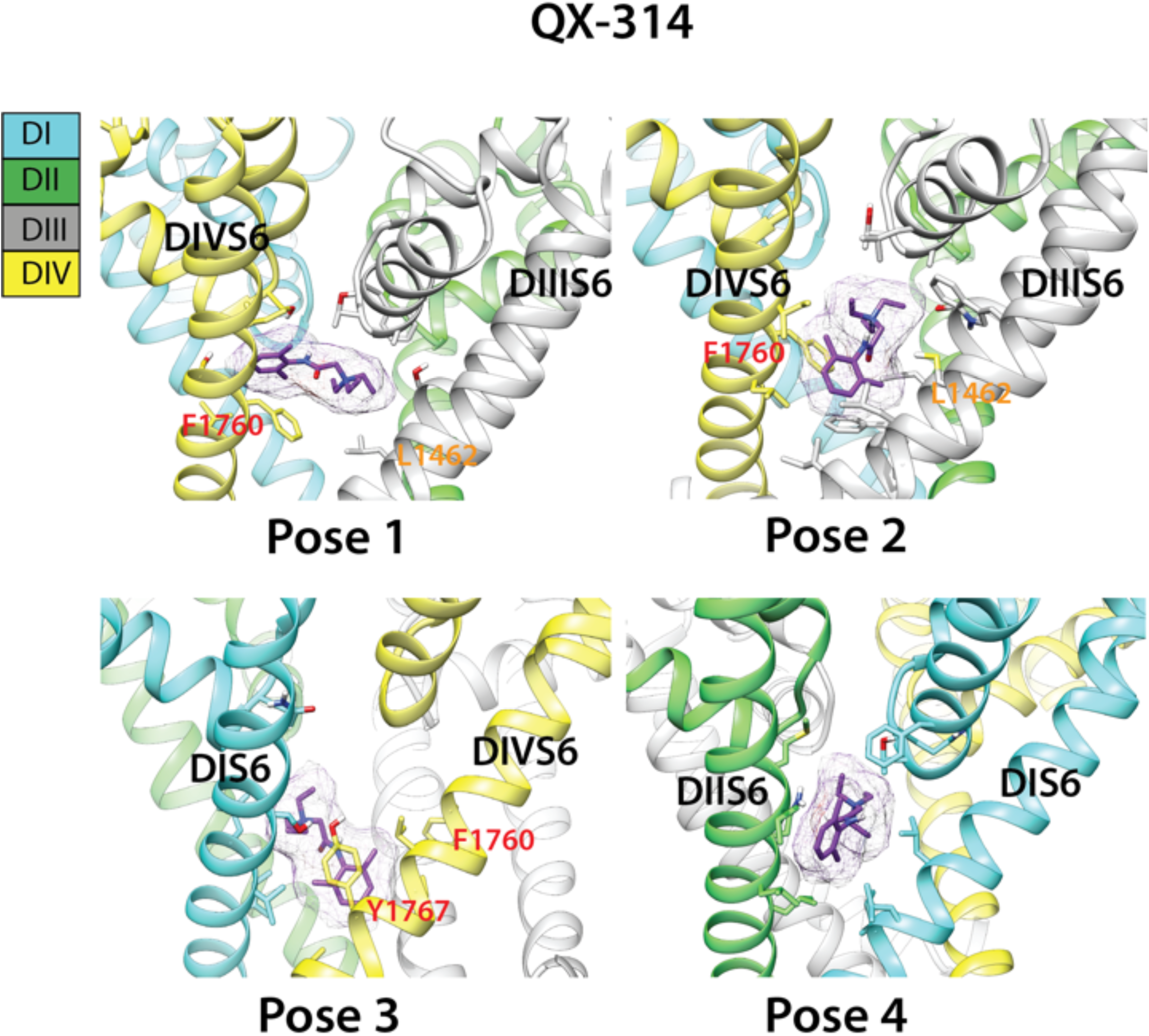
Top binding poses of QX-314 interaction with Rosetta model of hNav1.5 channel. Domain I is colored in blue, domain II is colored in green, domain III is colored gray, and domain IV is colored yellow. hNav1.5 residues forming interactions with lidocaine are shown in stick representation and labeled. QX-314 is shown in stick and surface representation and colored purple.

**Figure 2 - figure supplement 4.**
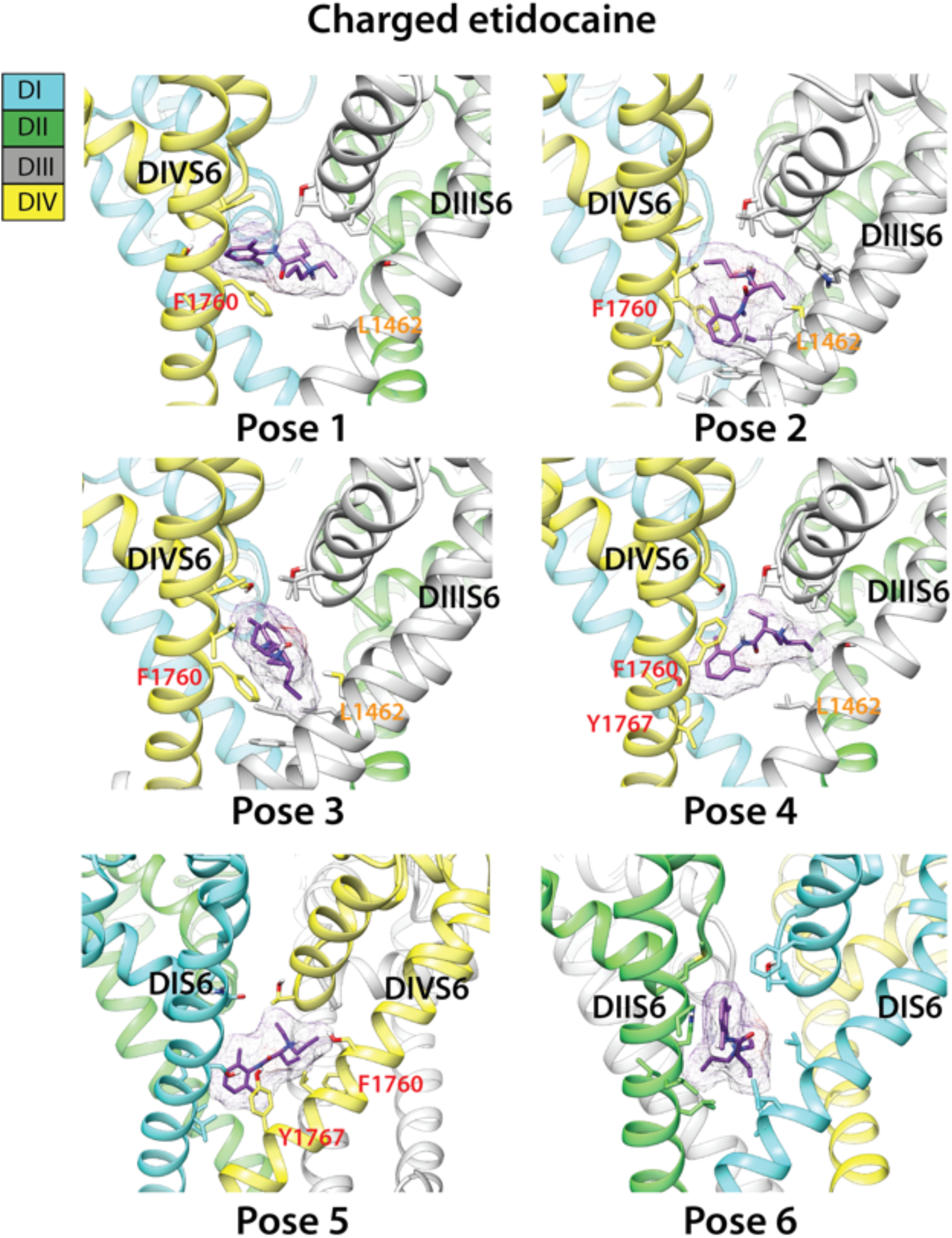
Top binding poses of charged etidocaine interaction with Rosetta model of hNav1.5 channel. Domain I is colored in blue, domain II is colored in green, domain III is colored gray, and domain IV is colored yellow. hNav1.5 residues forming interactions with lidocaine are shown in stick representation and labeled. Etidocaine is shown in stick and surface representation and colored purple.

**Figure 3 - figure supplement 1.**
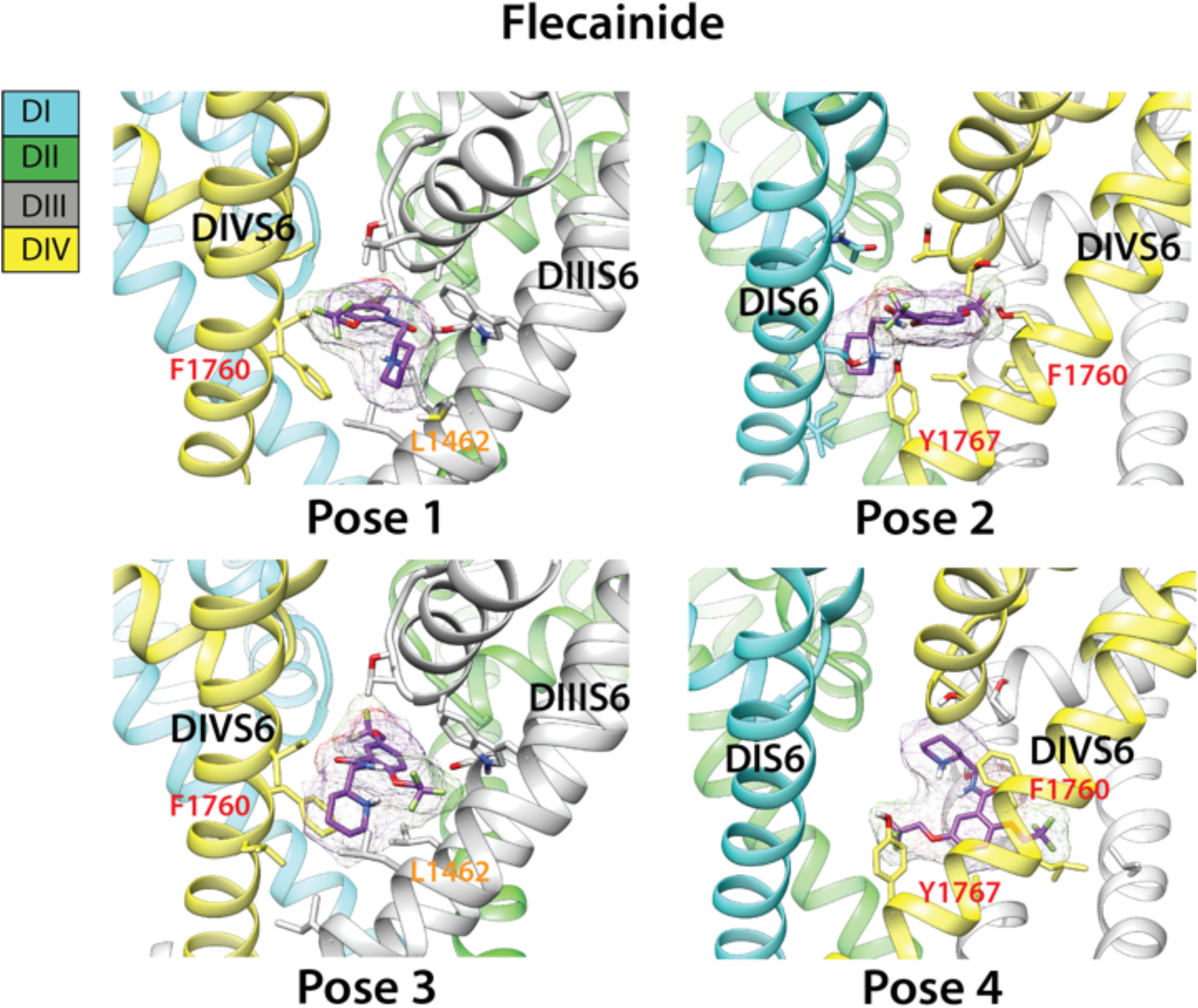
Top binding poses of flecainide interaction with Rosetta model of hNav1.5 channel. Domain I is colored in blue, domain II is colored in green, domain III is colored gray, and domain IV is colored yellow. hNav1.5 residues forming interactions with lidocaine are shown in stick representation and labeled. Flecainide is shown in stick and surface representation and colored purple.

**Figure 3 - figure supplement 2.**
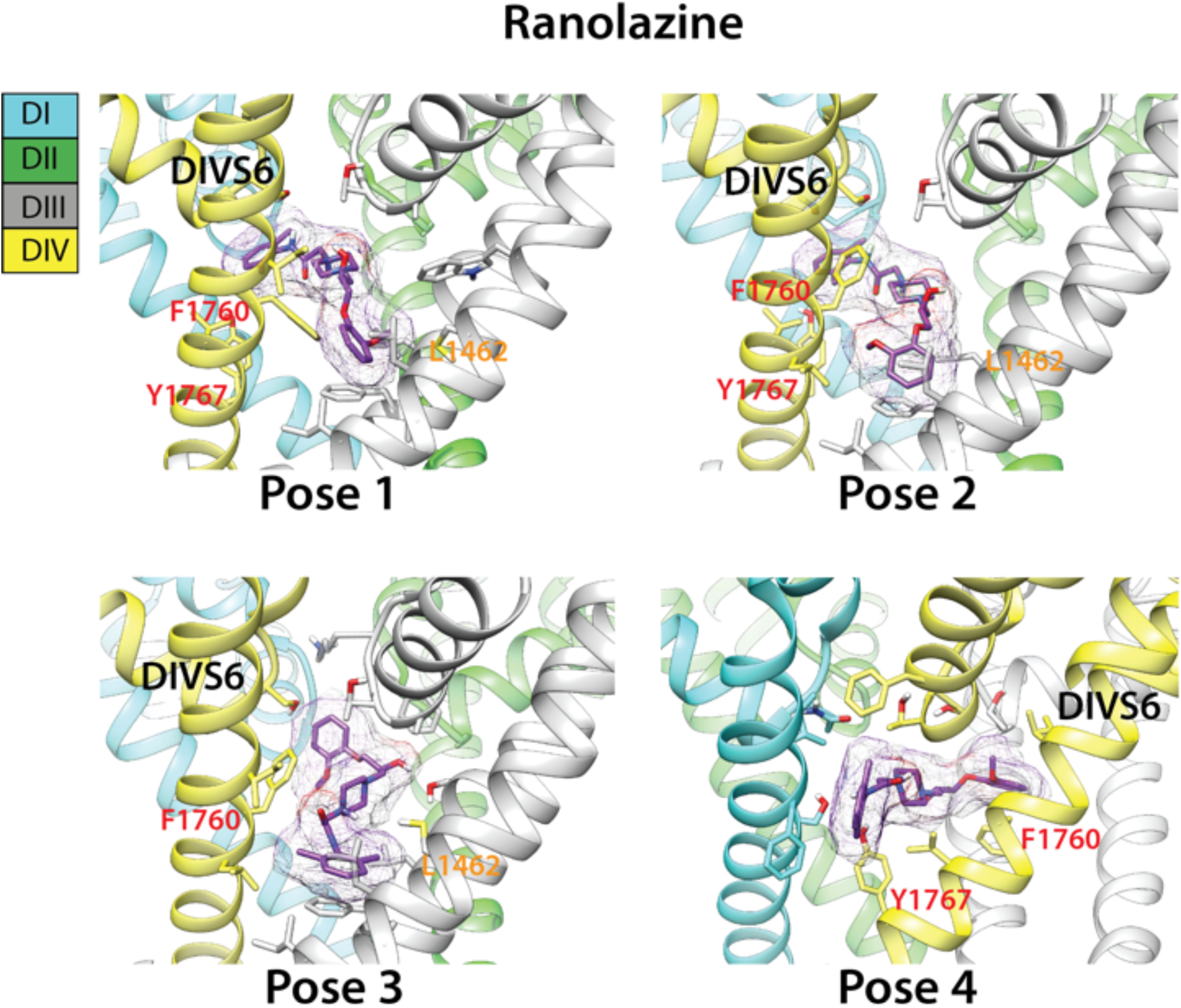
Top binding poses of ranolazine interaction with Rosetta model of hNav1.5 channel. Domain I is colored in blue, domain II is colored in green, domain III is colored gray, and domain IV is colored yellow. hNav1.5 residues forming interactions with lidocaine are shown in stick representation and labeled. Ranolazine is shown in stick and surface representation and colored purple.

**Figure 4 - figure supplement 1.**
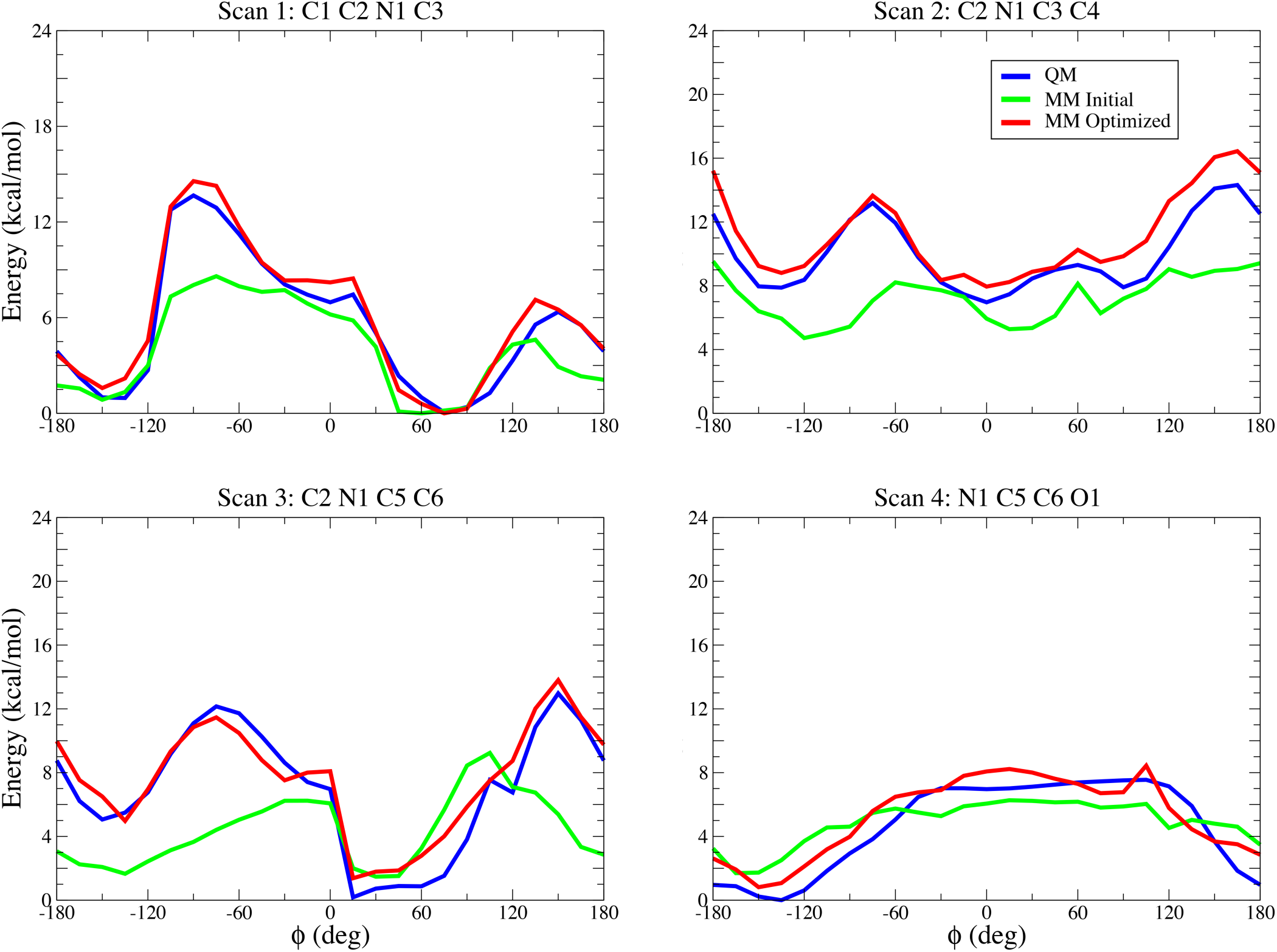
Gas-phase torsional energy profiles for neutral lidocaine (LID0) from quantum mechanical (QM), initial and optimized molecular mechanics (MM) calculations. Atom names correspond to ones in topology and parameter files.

**Figure 4 - figure supplement 2.**
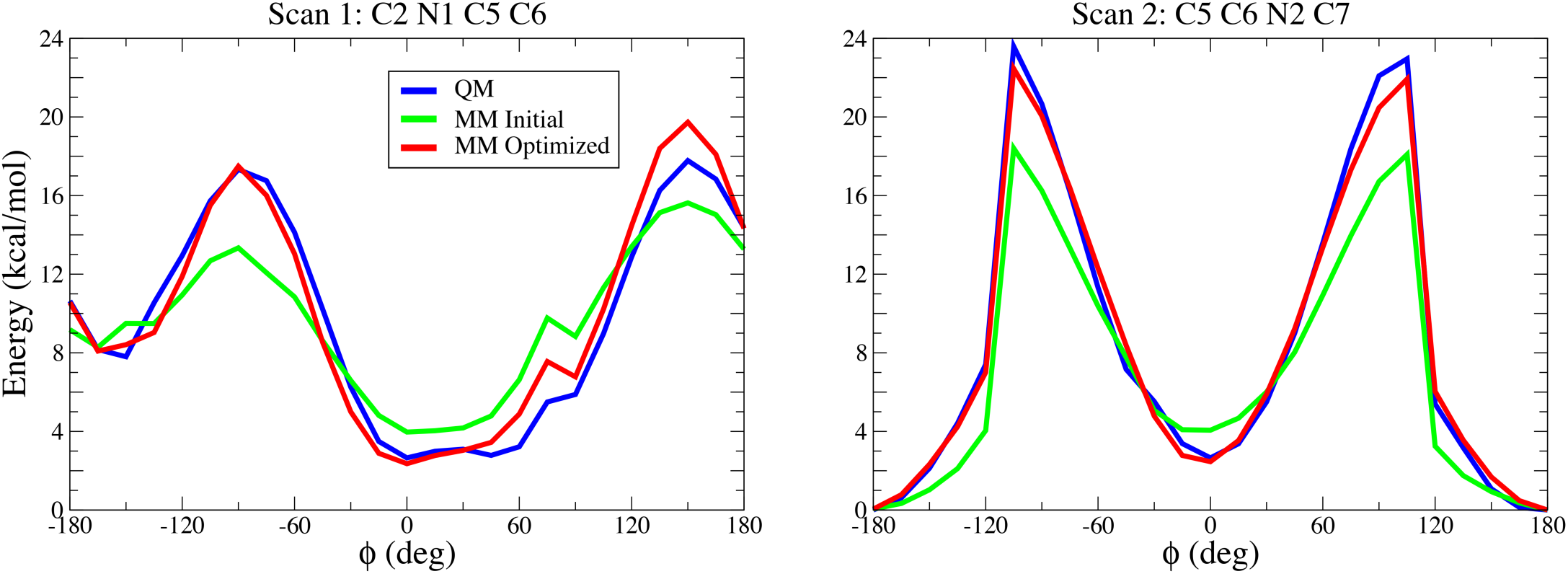
Gas-phase torsional energy profiles for charged lidocaine (LID1) from quantum mechanical (QM), initial and optimized molecular mechanics (MM) calculations. Atom names correspond to ones in topology and parameter files.

**Figure 4 - figure supplement 3.**
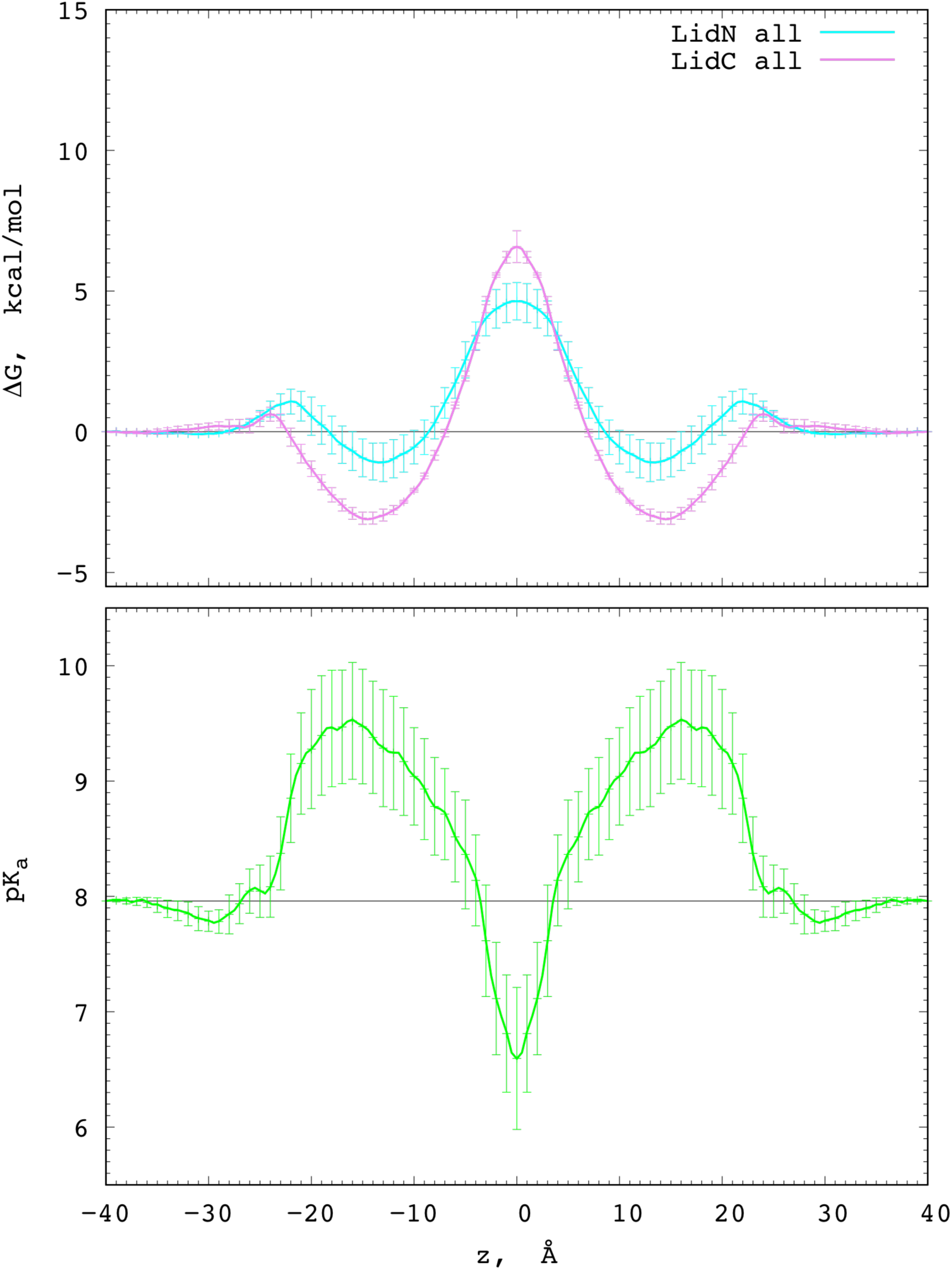
Charged and neutral lidocaine translocation across a POPC membrane. PMF profiles for POPC membrane crossing neutral (cyan) and charged (magenta) drug (top) and corresponding pKa profile (bottom). Error bars computed as a measure of asymmetry.

## Appendix 1

### Appendix S1. Charged lidocaine (LID1) optimized CHARMM force field topology and parameter files

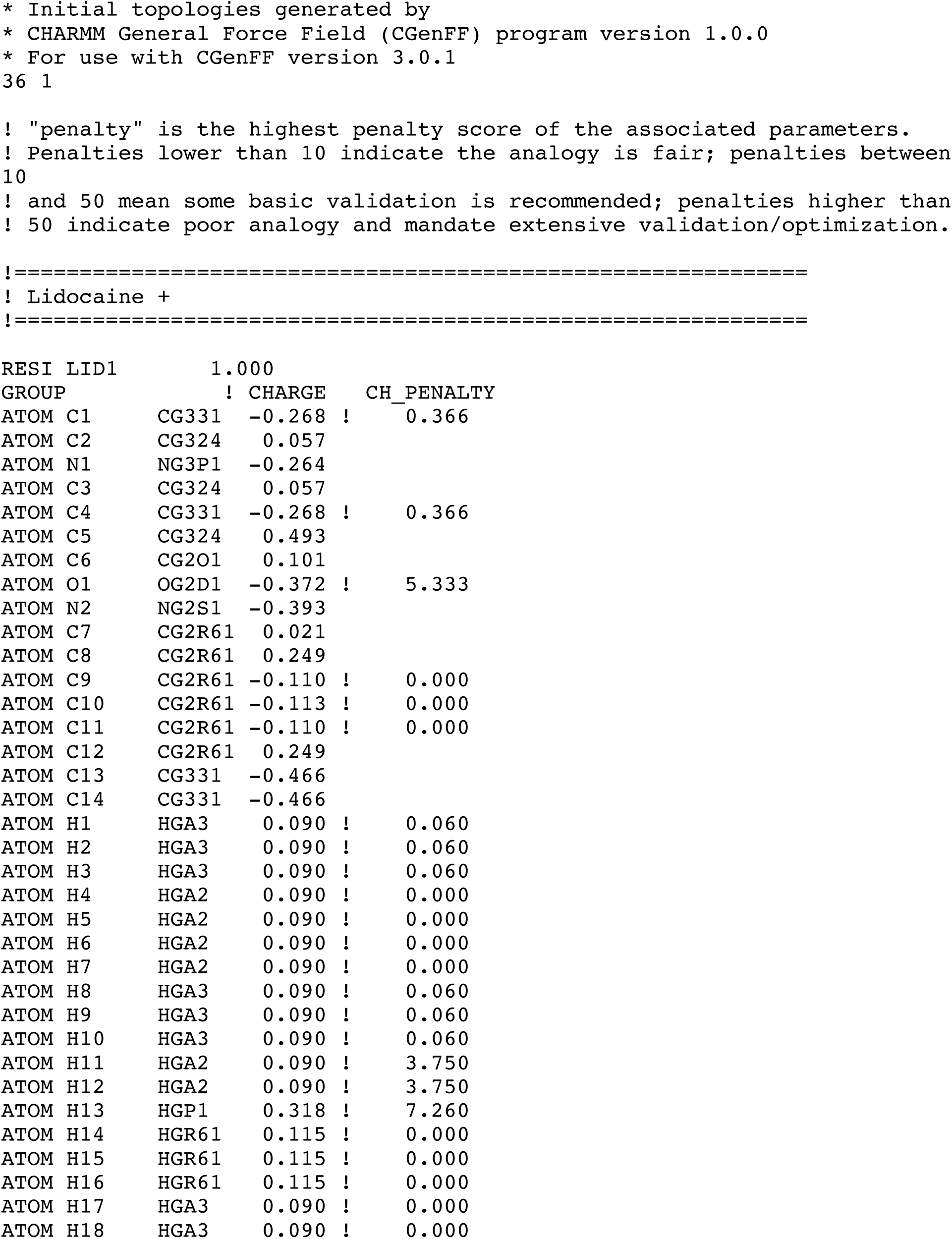
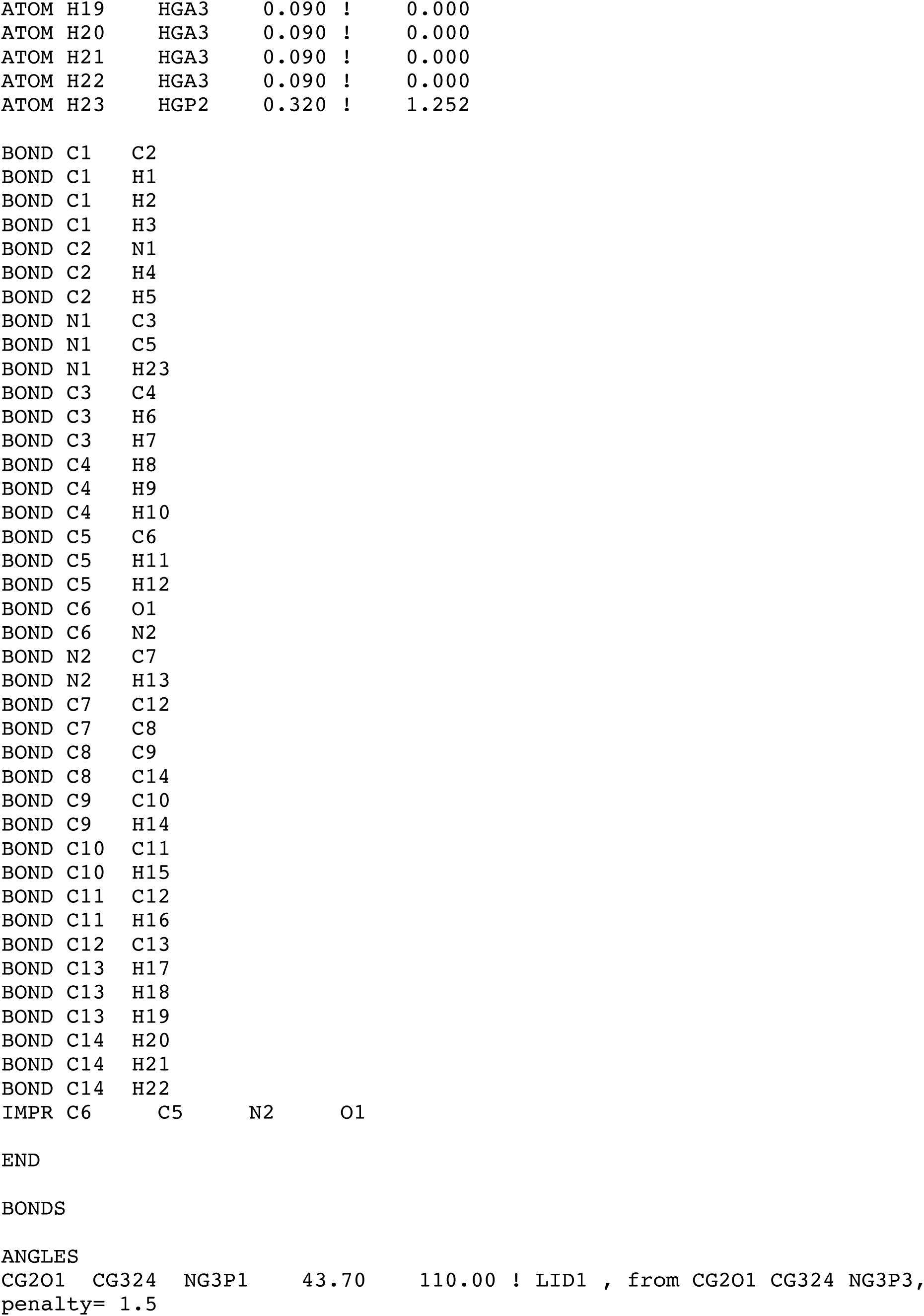
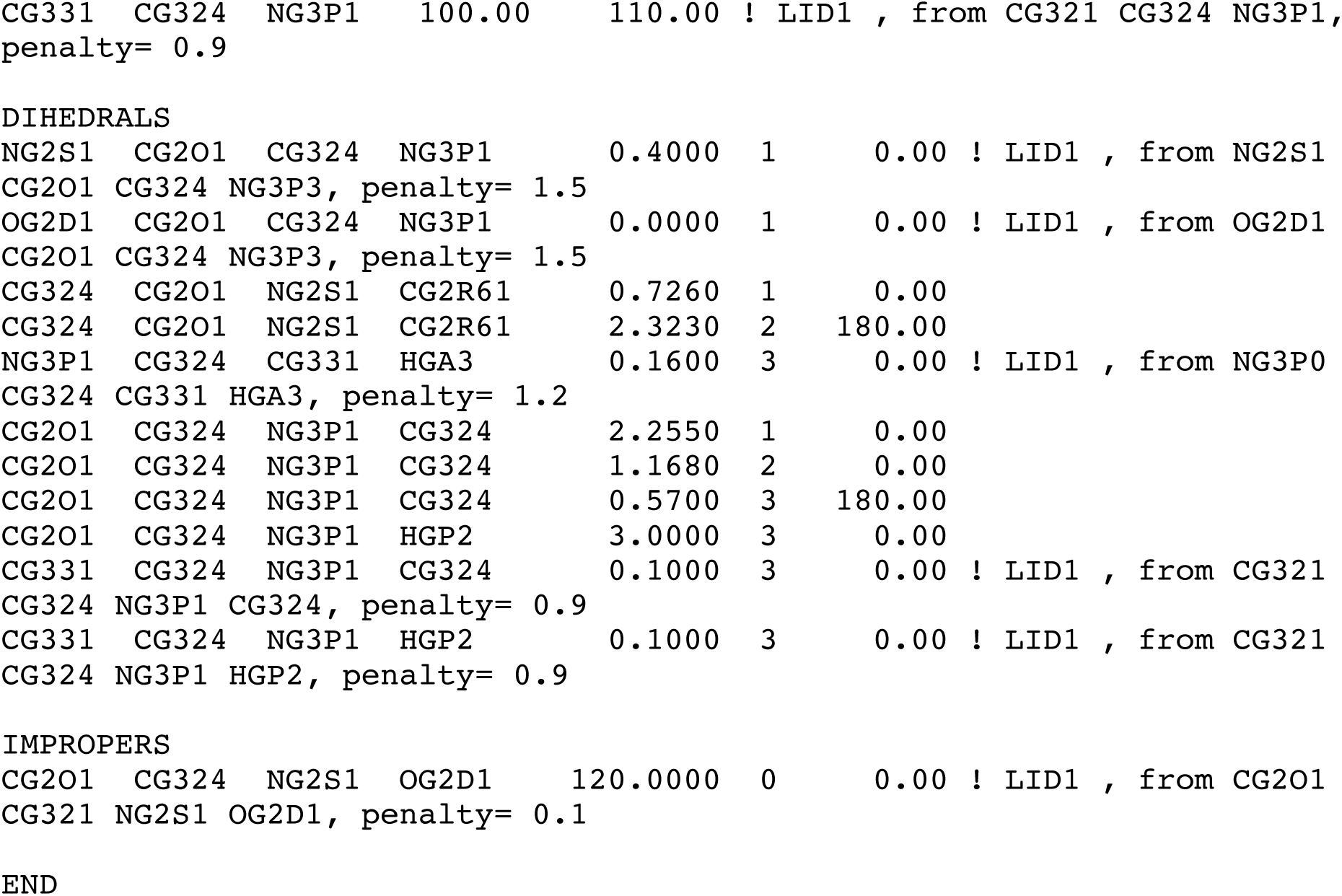

### Appendix S1. Neutral lidocaine (LID0) optimized CHARMM force field topology and parameter files

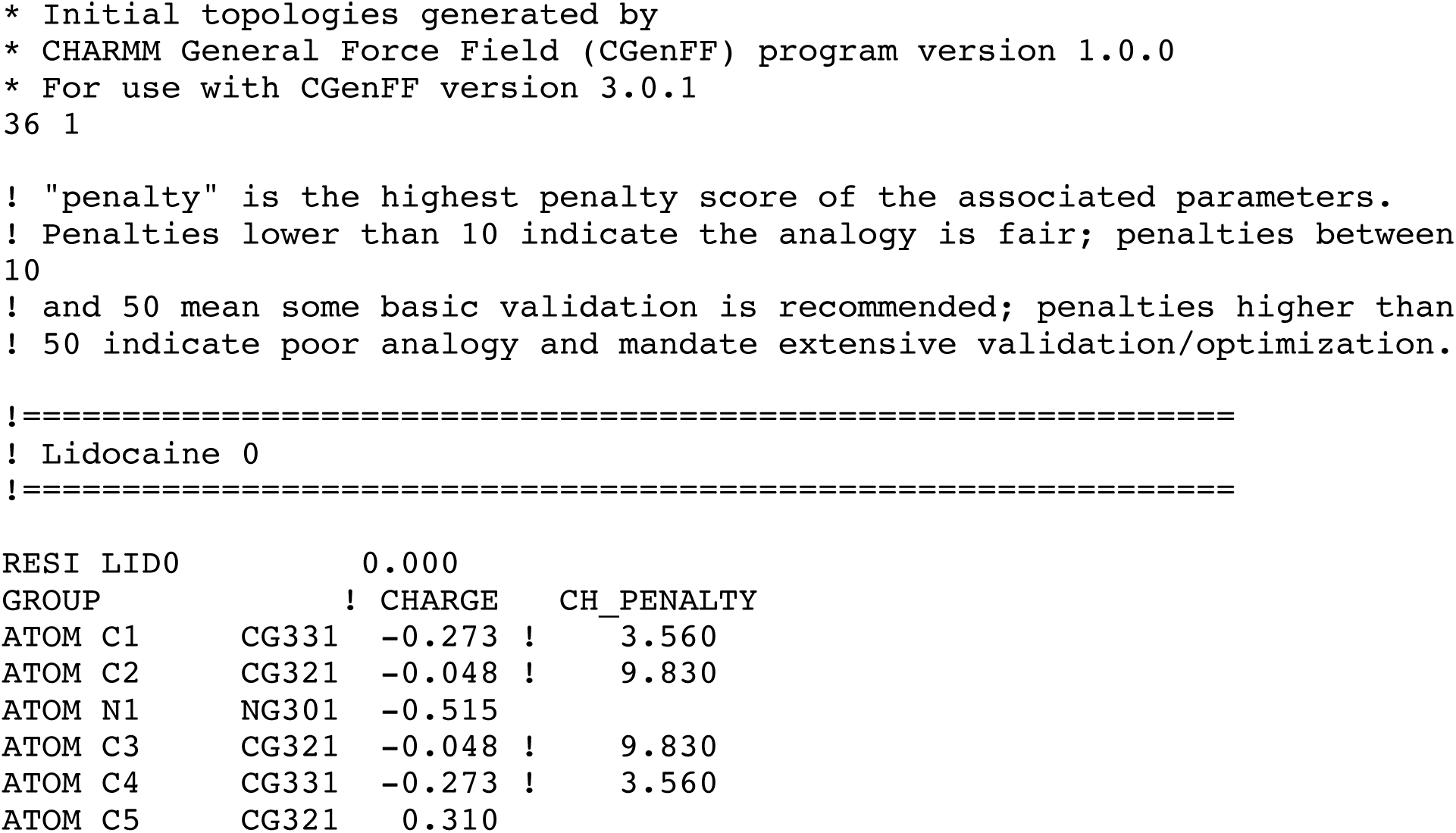
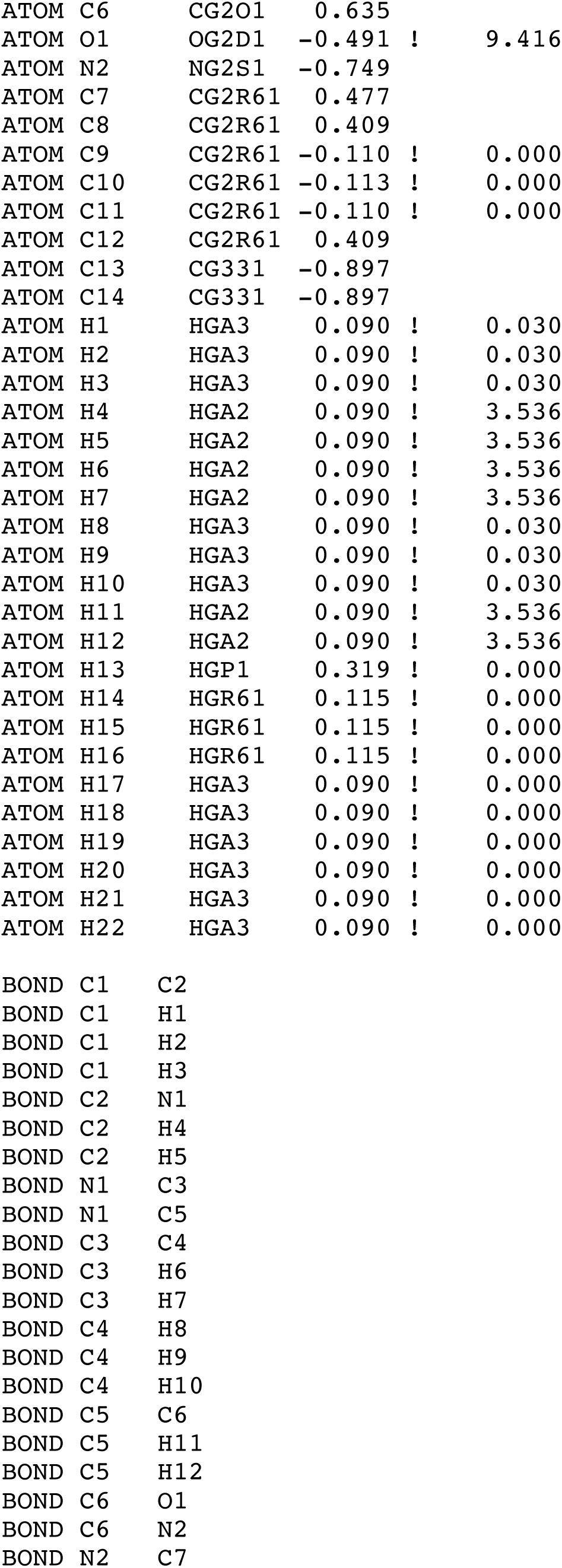
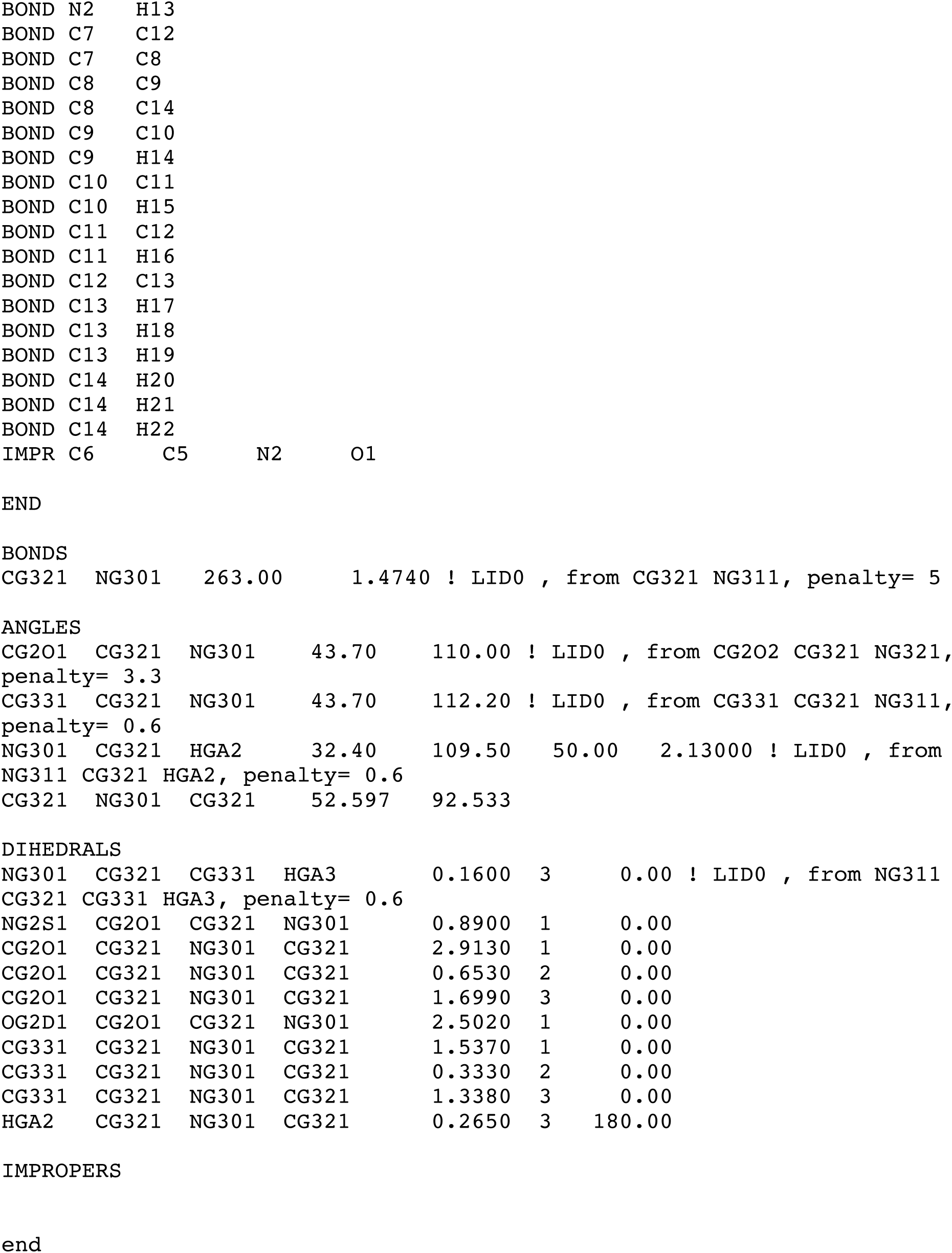

### Appendix S2. RosettaLigand docking scripts

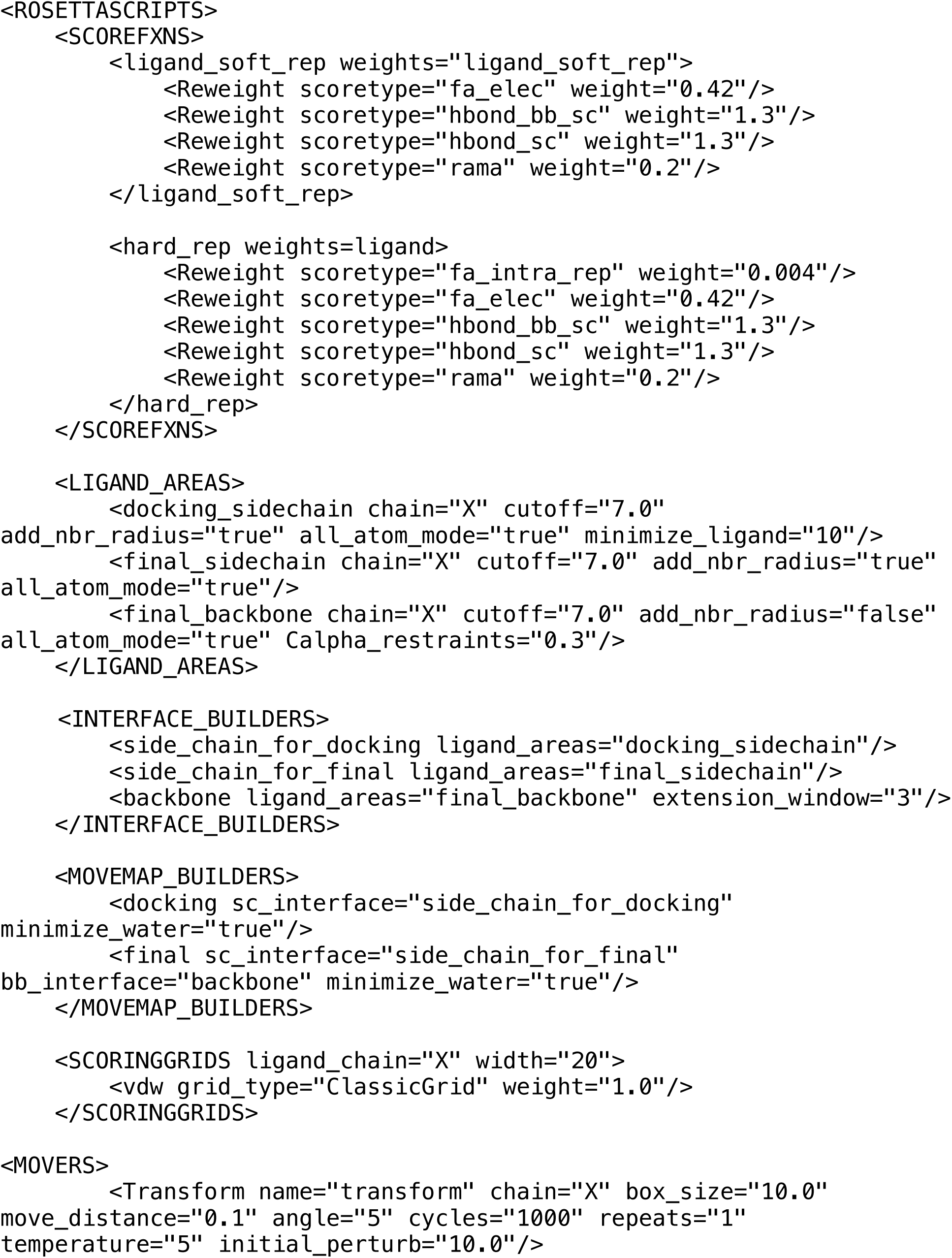
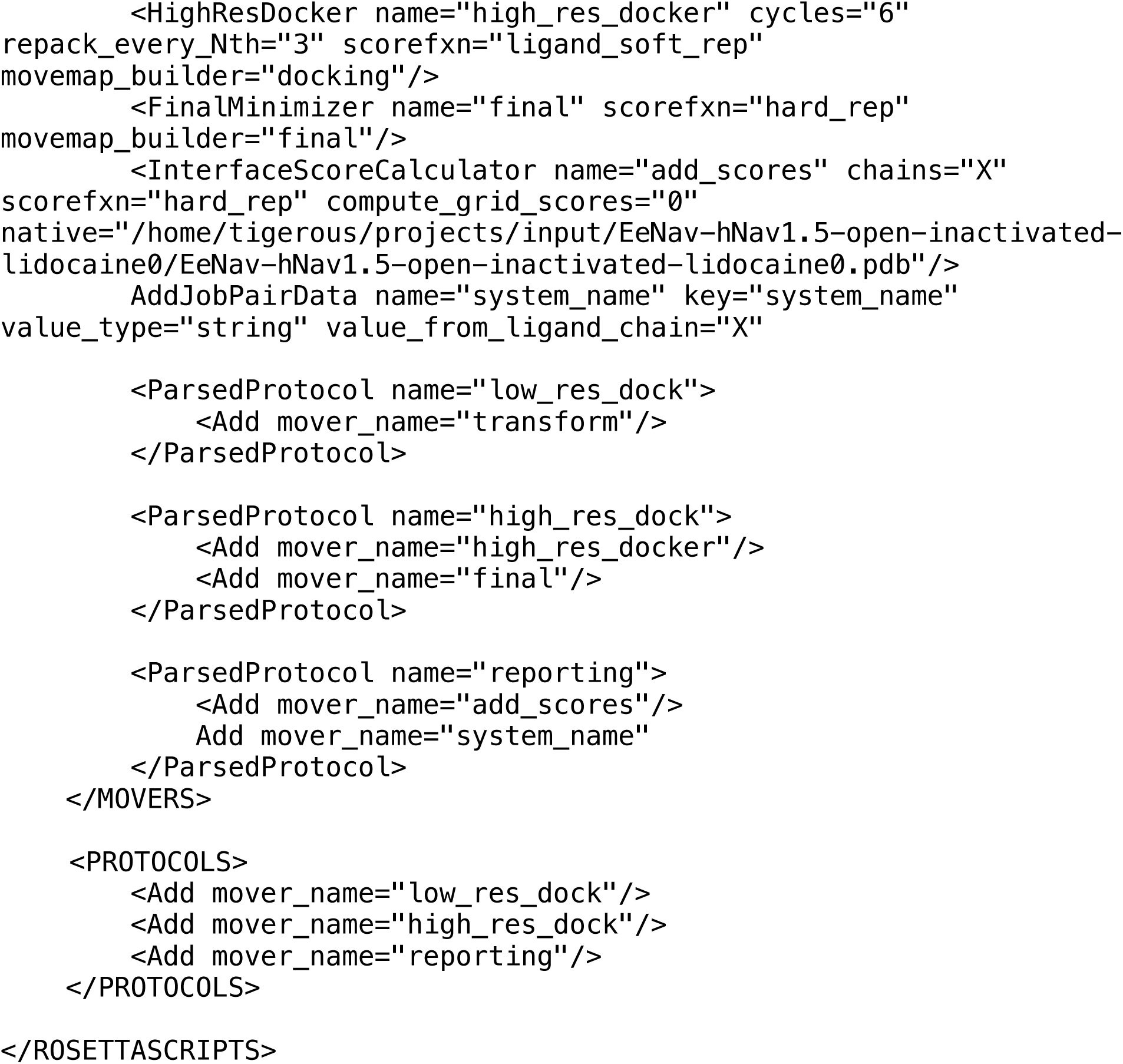

### Appendix S2. RosettaLigand docking flags

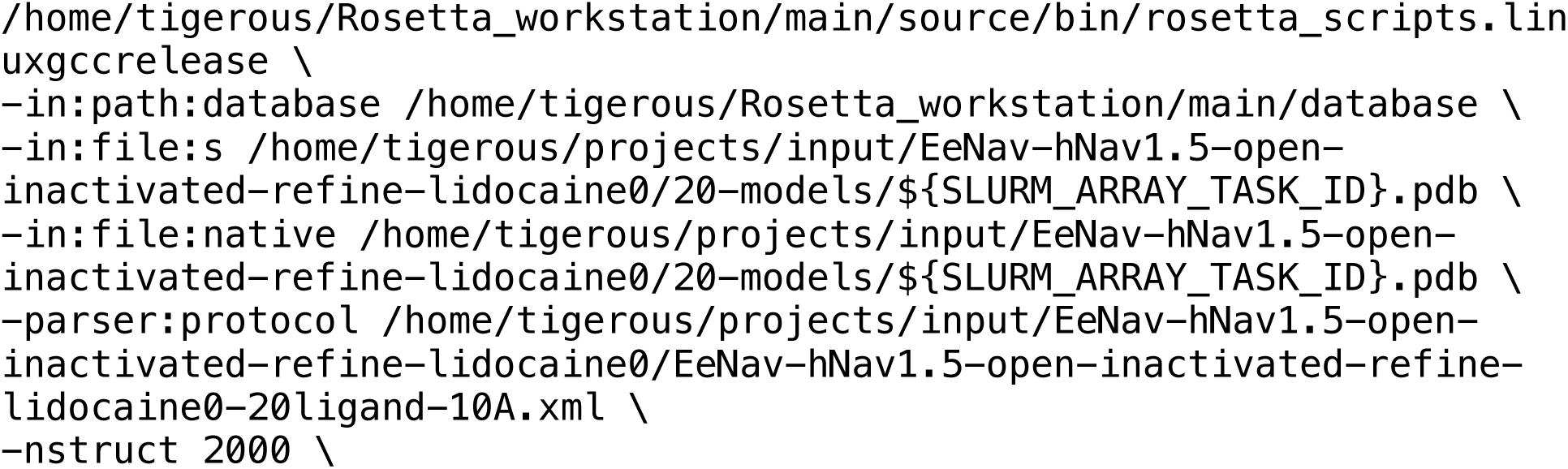
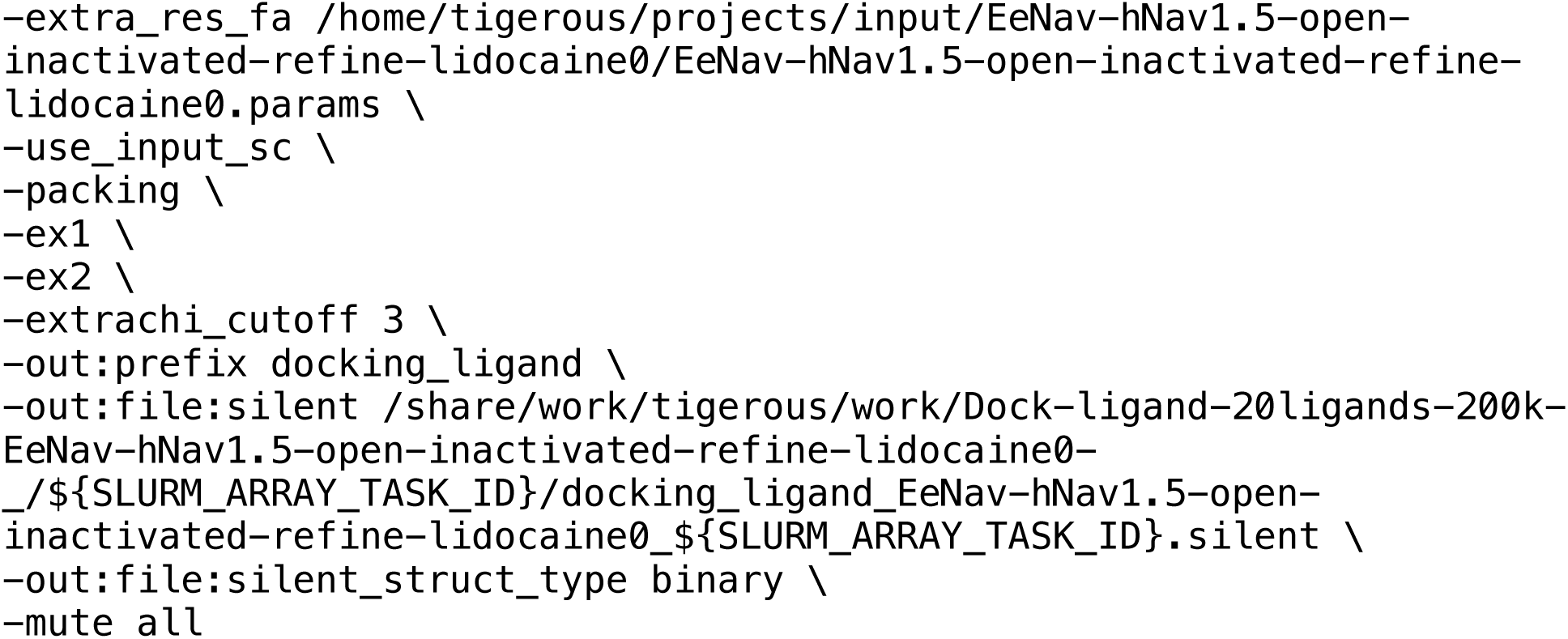

**Table S1.** Partial atomic charges for charged (LID1) and neutral (LID0) lidocaine models. (Optimized charge values are shown by asterisk)

|  | LID1 |  | LID0 |
| --- | --- | --- | --- |
| C1 | -0.268 | C1 | -0.273 |
| C2 * | 0.057 | C2 | -0.048 |
| N1 * | -0.264 | N1 * | -0.515 |
| C3 * | 0.057 | C3 | -0.048 |
| C4 | -0.268 | C4 | -0.273 |
| C5 * | 0.493 | C5 * | 0.31 |
| C6 * | 0.101 | C6 * | 0.635 |
| O1 | -0.372 | O1 | -0.491 |
| N2 * | -0.393 | N2 * | -0.749 |
| C7 * | 0.021 | C7 * | 0.477 |
| C8 * | 0.249 | C8 * | 0.409 |
| C9 | -0.11 | C9 | -0.11 |
| C10 | -0.113 | C10 | -0.113 |
| C11 | -0.11 | C11 | -0.11 |
| C12* | 0.249 | C12* | 0.409 |
| C13* | -0.466 | C13* | -0.897 |
| C14* | -0.466 | C14* | -0.897 |
| H1 | 0.09 | H1 | 0.09 |
| H2 | 0.09 | H2 | 0.09 |
| H3 | 0.09 | H3 | 0.09 |
| H4 | 0.09 | H4 | 0.09 |
| H5 | 0.09 | H5 | 0.09 |
| H6 | 0.09 | H6 | 0.09 |
| H7 | 0.09 | H7 | 0.09 |
| H8 | 0.09 | H8 | 0.09 |
| H9 | 0.09 | H9 | 0.09 |
| H10 | 0.09 | H10 | 0.09 |
| H11 | 0.09 | H11 | 0.09 |
| H12 | 0.09 | H12 | 0.09 |
| H13 | 0.318 | H13 | 0.319 |
| H14 | 0.115 | H14 | 0.115 |
| H15 | 0.115 | H15 | 0.115 |
| H16 | 0.115 | H16 | 0.115 |
| H17 | 0.09 | H17 | 0.09 |
| H18 | 0.09 | H18 | 0.09 |
| H19 | 0.09 | H19 | 0.09 |
| H20 | 0.09 | H20 | 0.09 |
| H21 | 0.09 | H21 | 0.09 |
| H22 | 0.09 | H22 | 0.09 |
| H23 | 0.32 |  |  |

**Table S2.**
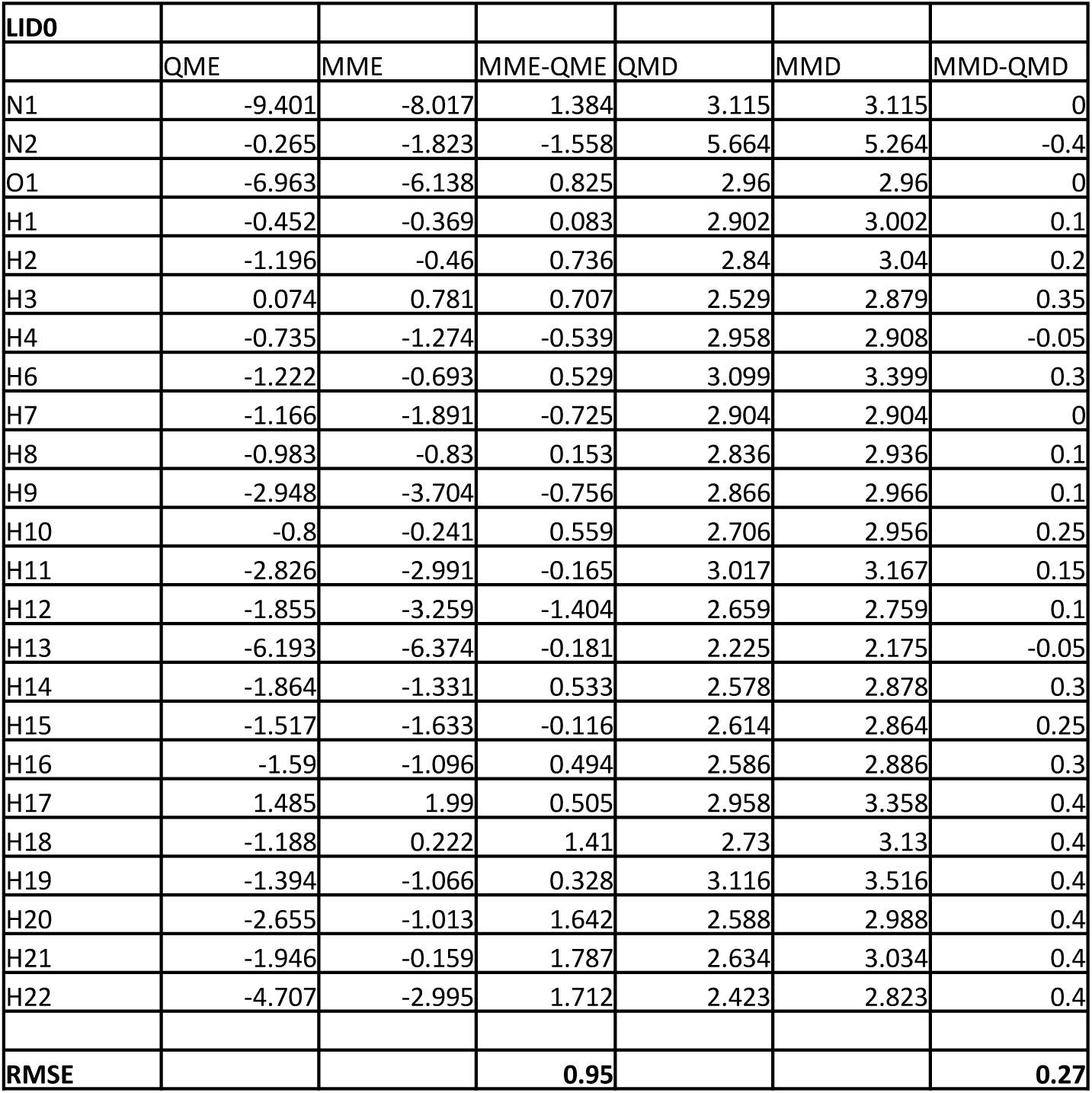
Gas-phase cationic lidocaine (LID1) - water interactions.

| LID0 |  |  |  |  |  |  |
| --- | --- | --- | --- | --- | --- | --- |
|  | QME | MME | MME-QME | QMD | MMD | MMD-QMD |
| N1 | -9.401 | -8.017 | 1.384 | 3.115 | 3.115 | 0 |
| N2 | -0.265 | -1.823 | -1.558 | 5.664 | 5.264 | -0.4 |
| O1 | -6.963 | -6.138 | 0.825 | 2.96 | 2.96 | 0 |
| H1 | -0.452 | -0.369 | 0.083 | 2.902 | 3.002 | 0.1 |
| H2 | -1.196 | -0.46 | 0.736 | 2.84 | 3.04 | 0.2 |
| H3 | 0.074 | 0.781 | 0.707 | 2.529 | 2.879 | 0.35 |
| H4 | -0.735 | -1.274 | -0.539 | 2.958 | 2.908 | -0.05 |
| H6 | -1.222 | -0.693 | 0.529 | 3.099 | 3.399 | 0.3 |
| H7 | -1.166 | -1.891 | -0.725 | 2.904 | 2.904 | 0 |
| H8 | -0.983 | -0.83 | 0.153 | 2.836 | 2.936 | 0.1 |
| H9 | -2.948 | -3.704 | -0.756 | 2.866 | 2.966 | 0.1 |
| H10 | -0.8 | -0.241 | 0.559 | 2.706 | 2.956 | 0.25 |
| H11 | -2.826 | -2.991 | -0.165 | 3.017 | 3.167 | 0.15 |
| H12 | -1.855 | -3.259 | -1.404 | 2.659 | 2.759 | 0.1 |
| H13 | -6.193 | -6.374 | -0.181 | 2.225 | 2.175 | -0.05 |
| H14 | -1.864 | -1.331 | 0.533 | 2.578 | 2.878 | 0.3 |
| H15 | -1.517 | -1.633 | -0.116 | 2.614 | 2.864 | 0.25 |
| H16 | -1.59 | -1.096 | 0.494 | 2.586 | 2.886 | 0.3 |
| H17 | 1.485 | 1.99 | 0.505 | 2.958 | 3.358 | 0.4 |
| H18 | -1.188 | 0.222 | 1.41 | 2.73 | 3.13 | 0.4 |
| H19 | -1.394 | -1.066 | 0.328 | 3.116 | 3.516 | 0.4 |
| H20 | -2.655 | -1.013 | 1.642 | 2.588 | 2.988 | 0.4 |
| H21 | -1.946 | -0.159 | 1.787 | 2.634 | 3.034 | 0.4 |
| H22 | -4.707 | -2.995 | 1.712 | 2.423 | 2.823 | 0.4 |
| RMSE |  |  | 0.95 |  |  | 0.27 |

**Table S3.**
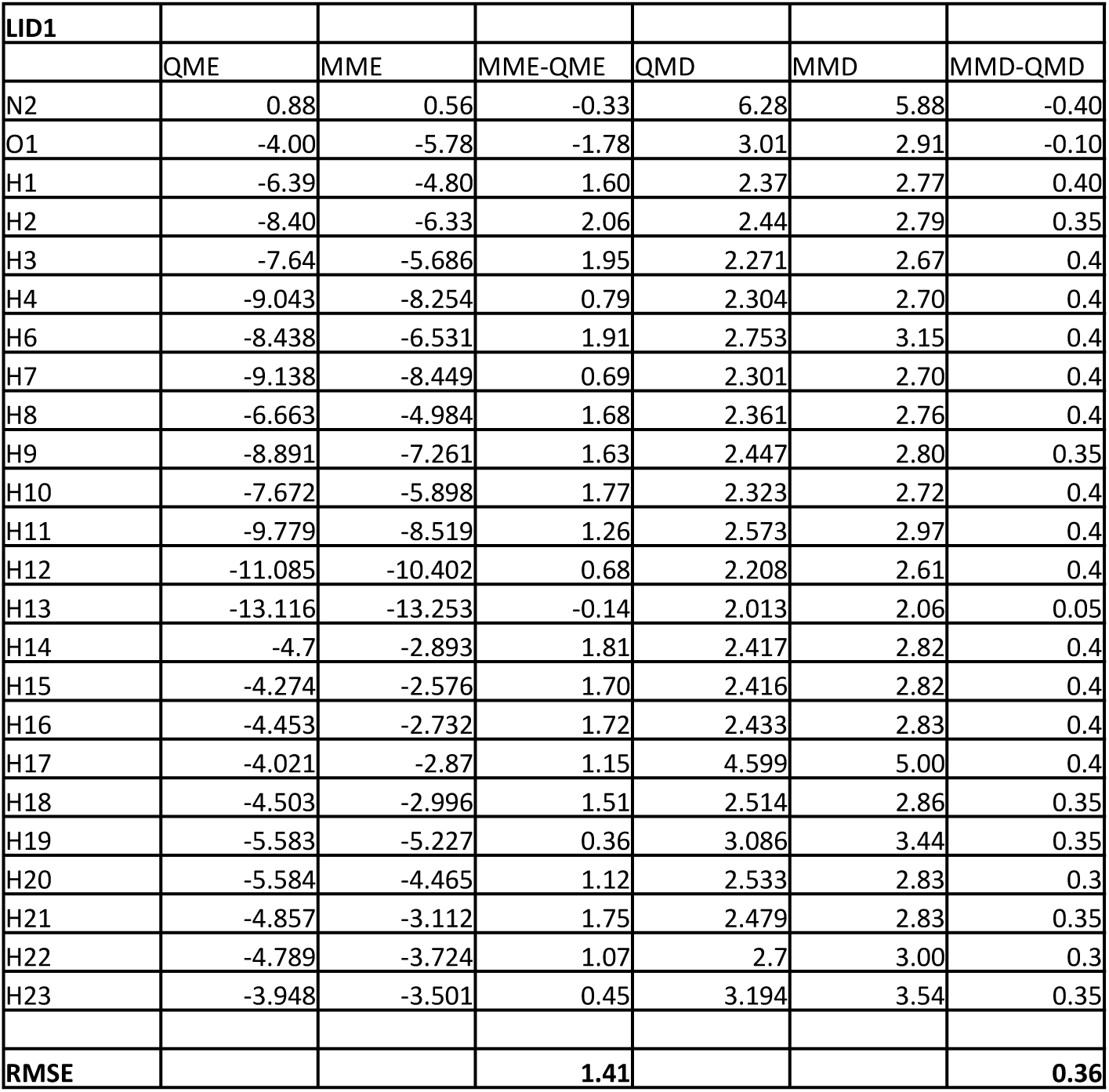
Gas-phase neutral lidocaine (LID0) - water interactions.

| LID1 |  |  |  |  |  |  |
| --- | --- | --- | --- | --- | --- | --- |
|  | QME | MME | MME-QME | QMD | MMD | MMD-QMD |
| N2 | 0.88 | 0.56 | -0.33 | 6.28 | 5.88 | -0.40 |
| O1 | -4.00 | -5.78 | -1.78 | 3.01 | 2.91 | -0.10 |
| H1 | -6.39 | -4.80 | 1.60 | 2.37 | 2.77 | 0.40 |
| H2 | -8.40 | -6.33 | 2.06 | 2.44 | 2.79 | 0.35 |
| H3 | -7.64 | -5.686 | 1.95 | 2.271 | 2.67 | 0.4 |
| H4 | -9.043 | -8.254 | 0.79 | 2.304 | 2.70 | 0.4 |
| H6 | -8.438 | -6.531 | 1.91 | 2.753 | 3.15 | 0.4 |
| H7 | -9.138 | -8.449 | 0.69 | 2.301 | 2.70 | 0.4 |
| H8 | -6.663 | -4.984 | 1.68 | 2.361 | 2.76 | 0.4 |
| H9 | -8.891 | -7.261 | 1.63 | 2.447 | 2.80 | 0.35 |
| H10 | -7.672 | -5.898 | 1.77 | 2.323 | 2.72 | 0.4 |
| H11 | -9.779 | -8.519 | 1.26 | 2.573 | 2.97 | 0.4 |
| H12 | -11.085 | -10.402 | 0.68 | 2.208 | 2.61 | 0.4 |
| H13 | -13.116 | -13.253 | -0.14 | 2.013 | 2.06 | 0.05 |
| H14 | -4.7 | -2.893 | 1.81 | 2.417 | 2.82 | 0.4 |
| H15 | -4.274 | -2.576 | 1.70 | 2.416 | 2.82 | 0.4 |
| H16 | -4.453 | -2.732 | 1.72 | 2.433 | 2.83 | 0.4 |
| H17 | -4.021 | -2.87 | 1.15 | 4.599 | 5.00 | 0.4 |
| H18 | -4.503 | -2.996 | 1.51 | 2.514 | 2.86 | 0.35 |
| H19 | -5.583 | -5.227 | 0.36 | 3.086 | 3.44 | 0.35 |
| H20 | -5.584 | -4.465 | 1.12 | 2.533 | 2.83 | 0.3 |
| H21 | -4.857 | -3.112 | 1.75 | 2.479 | 2.83 | 0.35 |
| H22 | -4.789 | -3.724 | 1.07 | 2.7 | 3.00 | 0.3 |
| H23 | -3.948 | -3.501 | 0.45 | 3.194 | 3.54 | 0.35 |
| RMSE |  |  | 1.41 |  |  | 0.36 |

